# α-carboxysome formation is mediated by the multivalent and disordered protein CsoS2

**DOI:** 10.1101/708164

**Authors:** Luke M. Oltrogge, Thawatchai Chaijarasphong, Allen W. Chen, Eric R. Bolin, Susan Marqusee, David F. Savage

**Author notes:** Department of Biotechnology, Faculty of Science, Mahidol University, Rama VI Rd., Bangkok 10400, Thailand.

## Abstract

Carboxysomes are bacterial microcompartments that function as the centerpiece of the bacterial CO_2_-concentrating mechanism, feeding high concentrations of CO_2_ to the enzyme Rubisco for fixation. The carboxysome self-assembles from thousands of individual proteins into icosahedral-like particles with a dense enzyme cargo encapsulated within a proteinaceous shell. In the case of the α-carboxysome, there is little molecular insight into protein-protein interactions which drive the assembly process. Here we show that the N-terminus of CsoS2, an intrinsically disordered protein found in the α-carboxysome, possesses a repeated peptide sequence that binds Rubisco. X-ray structural analysis of the peptide bound to Rubisco reveals a series of conserved electrostatic interactions that are only made with properly assembled hexadecameric Rubisco. Although biophysical measurements indicate this single interaction is weak, its implicit multivalency induces high-affinity binding through avidity. Taken together, our results indicate CsoS2 acts as an interaction hub to condense Rubisco and enable efficient α-carboxysome formation.

## Introduction

Many carbon-assimilating bacteria possess CO_2_-concentrating mechanisms (CCMs) to facilitate carbon fixation by the enzyme Rubisco.^1^ The centerpiece of the CCM is the carboxysome, a large protein complex which encapsulates Rubisco and carbonic anhydrase and is thought to produce locally high concentrations of CO_2_.^2,3^ The carboxysome is a large (100-400 nm diameter) and composite (∼10 different protomers) structure comprising both a virus-like protein shell and cargo enzymes.^4–6^ Moreover, carboxysome formation requires thousands of individual proteins to accurately self-assemble.^7–9^ How this mesoscopic complex, with linear dimensions roughly ten-fold larger than any of its individual components, assembles with high structural and compositional fidelity remains unknown.

Carboxysomes occur in two distinct evolutionary lineages, α and β, that are functionally and morphologically similar.^4,10,11^ Both enclose a dense enzymatic cargo of Rubisco and carbonic anhydrase inside the icosahedral shell composed of hexameric and pentameric proteins. One or more scaffolding proteins serve as interaction hubs, mediating the associations among the various components.^4^

Although the α-carboxysome was the first to be identified and characterized,^12^ the β-carboxysome assembly process is better understood. Two proteins, CcmM and CcmN, act in tandem as the scaffold in a hierarchical set of interactions to bridge shell with cargo.^4,13^ An amphipathic encapsulation peptide on CcmN anchors to CcmK, a hexameric shell protein.^14^ CcmN also binds to CcmM, a scaffolding protein which contains three to five tandem repeats of a Rubisco small subunit like (SSUL) module separated by disordered linkers. SSUL repeats then interact with Rubisco.^15–18^ Contrary to expectations based on sequence homology, SSULs do not displace the Rubisco small subunit but bind across the interface of two L_2_ dimers and a small subunit.^17^

The assembly of α-carboxysomes—the predominant form among oceanic cyanobacteria and autotrophic proteobacteria—is, to date, more opaque. One unique component of the α-carboxysome is CsoS2, a large (∼900 residues) intrinsically disordered protein (IDP), which, unlike CcmM or CcmN, contains no recognizable domains.^19,20^ CsoS2 is indispensable for carboxysome assembly and thus hypothesized to be a potential scaffolding protein. Knock-outs in the α-carboxysome model organism *Halothiobacillus neapolitanus* produce high CO_2_-requiring phenotypes and result in no observable carboxysomes.^19,21^ Pulldown and native agarose gel-shift assays using purified protein have demonstrated that CsoS2 interacts with both Rubisco and CsoS1 hexameric shell proteins.^19,22–24^ The specific sites of interaction, however, have not been definitively determined nor is it clear how they collectively give rise to robust assembly.

Here, we show that a repeated peptide motif in the N-terminal domain of CsoS2 interacts with Rubisco to facilitate encapsulation into the carboxysome. Using a fusion of this peptide with Rubisco we obtained a structure of the binding site which revealed a predominantly electrostatic interaction interface mediated by highly conserved residues. This binding site lies at a conjunction of Rubisco subunits uniquely present in the complete L_8_S_8_ oligomer, thus ensuring the encapsulation of only the functional holoenzyme. Energetic characterization indicated that the individual peptide/Rubisco interaction is very weak and relies on the engagement of multiple binding sites to increase its interaction strength. Bioinformatic analysis and expression of CsoS2-truncated heterologous carboxysomes implicate the multivalency of this interaction as an essential feature of the assembly process. Our data suggest that CsoS2 acts as a protein interaction hub which gathers Rubisco to nascent carboxysome shell facets through branching low-affinity interactions that collectively give rise to efficient and robust cargo accumulation.

## Results

### CsoS2 interacts with Rubisco

We and others have demonstrated the essentiality of CsoS2 to α-carboxysome formation.^19,21^ This fact, in combination with CsoS2’s unique sequence characteristics,^20^ led us to consider whether it is the scaffolding protein driving assembly of the α-carboxysome. CsoS2 is a repetitive IDP.^19,25^ It can be divided into three major domains, the N-terminal domain (NTD), Middle region (MR), and C-terminal domain (CTD), based on sequence self-similarity of the repeated motifs contained therein.^19^ The full protein has a high PONDR-FIT disorder score^26^ throughout (average = 0.63, >0.5 predicts disordered) and is only predicted to possess secondary structure within the repeats of the NTD (hereafter generically referred to as the ‘N-peptide’ or specifically by numbers, e.g. N1 through N4; Fig. 1a).^27^ Circular dichroism (CD) spectra indicated that only the NTD has α-helical content (Fig. 1b). However, the repeat sequences in the NTD do not necessarily coincide with regions of greater predicted order. It is thus possible that the N-peptides are in dynamic equilibrium between helical and unstructured conformations.

**Figure 1.**
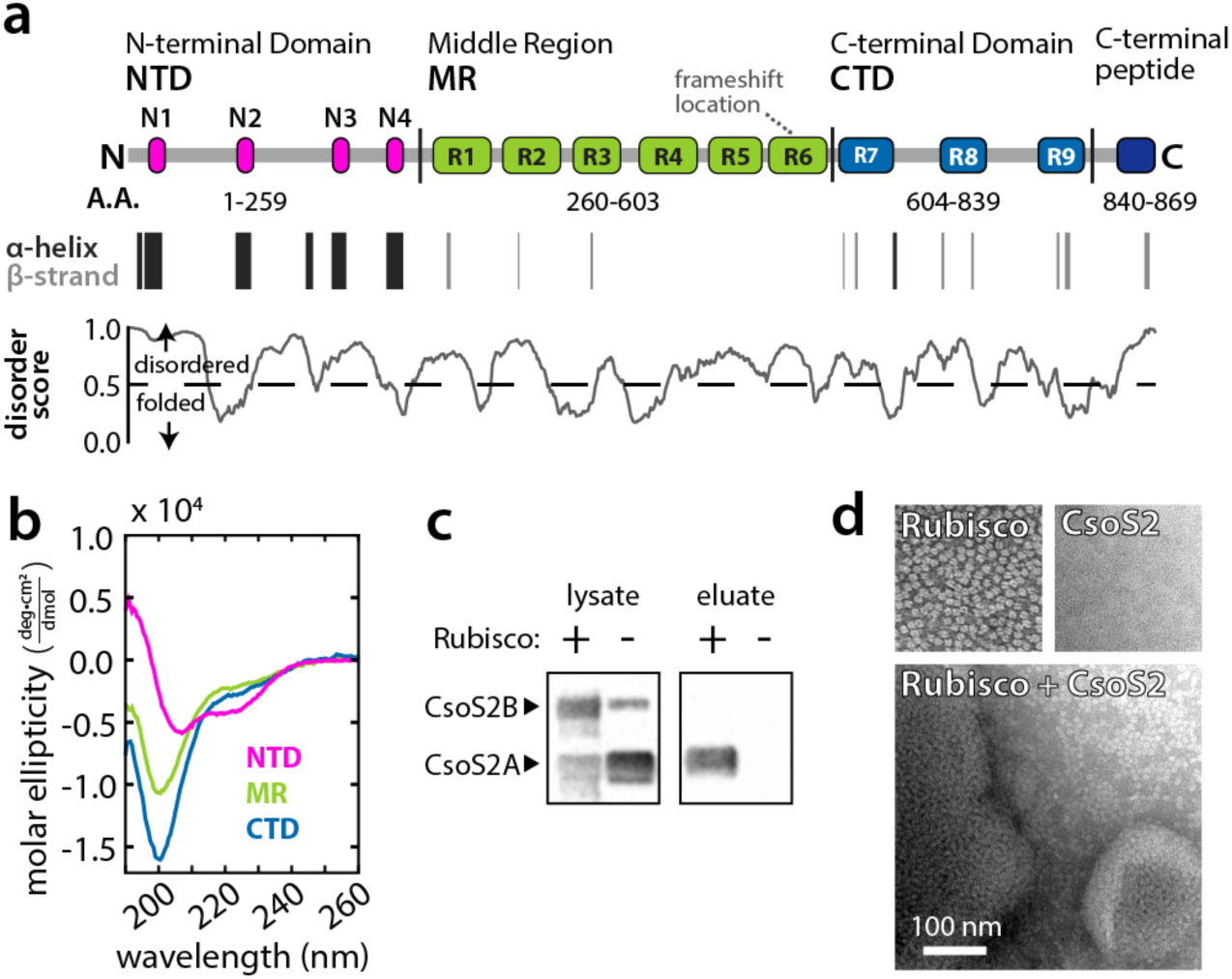
**a**, Repeat structure of *H. neapolitanus* CsoS2 with secondary structure prediction and disorder scores. “Frameshift location” indicates the site of a programmed ribosomal frameshift which results in expression of about 50% prematurely truncated protein (CsoS2A) and 50% full-length protein (CsoS2B).21 **b**, Circular dichroism spectra of each of the CsoS2 domains. **c**, Anti-His Western blot against His-tagged CsoS2 expressed +/-Strep-tagged Rubisco in the raw lysate and following Strep affinity purification (eluate). **d**, Negative stain TEM micrographs of purified Rubisco, CsoS2, and the aggregates observed when mixed.

Rubisco and CsoS2 together constitute a significant fraction of the cargo mass in purified carboxysomes and have complementary isoelectric points (5.9 and 9.1, respectively) suggesting a possible electrostatic association.^5^ We therefore tested whether these two proteins physically interact via pull-down analysis. As hypothesized, affinity purification of Strep-tagged Rubisco pulled down a 6xHis-tagged CsoS2 when visualized by anti-His Western blotting (Fig. 1c). This result pointed toward a direct interaction between CsoS2 and Rubisco and corroborated prior evidence.^19^ Furthermore, we observed dense aggregates of CsoS2 and Rubisco by transmission electron microscopy (TEM) when the two proteins were co-incubated (Fig. 1d).

### Repeated NTD motif binds Rubisco with low affinity

We next sought to identify the specific element of CsoS2 capable of interacting with Rubisco. This was carried out using bio-layer interferometry (BLI)—a label-free optical technique that monitors recruitment of a “prey” protein by a surface-immobilized “bait.”^28^ BLI analysis on CsoS2 and its various fragments revealed that binding activity resided in the NTD (Fig. 2a). IDPs often interact with their targets through short linear motifs^29,30^ and further analysis demonstrated that a single peptide derived from the consensus sequence of N1-N4, which we term N* (with sequence GRDLARARREALSQQGKAAV), was capable of interacting with Rubisco. A randomized sequence of N* (GRRKGLRAAGRALQVEQADSRA) did not bind (Fig. 2a,b), nor did any of the other conserved peptides from the MR or CTD (Fig. S2), suggesting that the interaction was indeed sequence specific and not, for example, due to generic charge-charge attraction.

**Figure 2.**
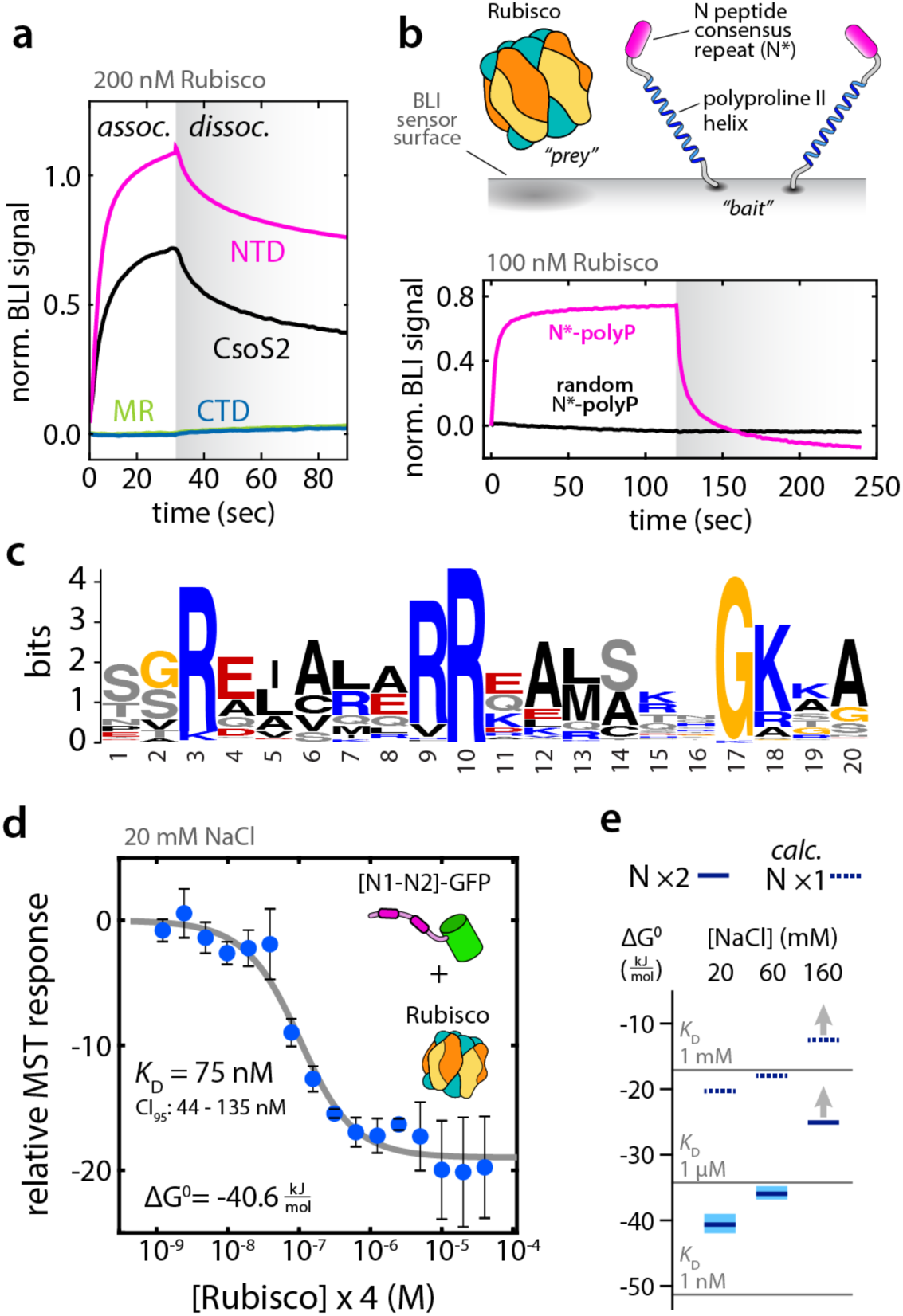
**a**, Bio-layer interferometry (BLI) Rubisco binding response normalized to the bait loading signal for full-length CsoS2 and each of the domains. **b**, Upper panel: schematic of the BLI sensor surface with the N*-peptide displayed on an extended polyproline II helix as the bait and Rubisco as the prey species. Lower panel: BLI response shows active binding of Rubisco by N* but not by a scrambled version. **c**, Weblogo conservation of the N-peptide motif calculated by MEME31 from 231 CsoS2 sequences which contained 901 N-peptide occurrences. **d**, Microscale thermophoresis (MST) binding isotherm with the first two *H. neapolitanus* CsoS2 N-peptide repeats fused to GFP, [N1-N2]-GFP, as the target and Rubisco as the ligand. The abscissa represents the concentration of binding sites for [N1-N2]-GFP, i.e. four per Rubisco. Error bars indicate +/-one standard deviation for measurements performed in triplicate. 95% confidence interval (CI_95_) estimated by bootstrap analysis. **e**, Standard free energies of binding for the reaction in (**d**) calculated from binding isotherms at 20, 60, and 160 mM NaCl. Solid dark blue lines are measured for [N1-N2]-GFP with light blue spanning the 95% confidence interval. Dashed blue lines are calculated estimates of the binding energy of a single repeat to a single Rubisco binding site. At 160 mM NaCl, no binding could be detected and the lines represent lower limits of the *K*_D_.

The interaction appeared to be driven by a specific sequence of positively charged residues. We analyzed a set of 231 CsoS2 sequences from α-cyanobacteria and proteobacteria with α-carboxysomes to identify the pan-species consensus N-peptide motif (Fig. 2c), recapitulating previous results.^19^ Notably, among the most highly conserved positions in the N-peptide motif are basic residues at positions 3, 9, 10, and 18, implying that the interaction likely has significant ionic character. R to A mutations were made for positions 3 and 10 in all of the four repeats in the NTD and entirely eliminated the binding in BLI (Fig. S3). Furthermore, a retrospective statistical examination of CsoS2 peptide array binding data from Cai et al.^19^ revealed a significant enrichment of Rubisco binding to peptides matching the N-peptide arginine motif (Fig. S7).

In principle, the binding energy between Rubisco and the N-peptide should be calculable from fitting the association and dissociation kinetics. However, due to the inherently high valency of the L_8_S_8_ Rubisco complex and the surface-induced avidity of neighboring bait proteins, it was difficult to obtain reliable fits to a simple binding model (Fig. S1). For this reason, the solution-phase technique microscale thermophoresis (MST) was used to measure binding in an alternative fashion. Unexpectedly, while the implied dissociation constants (*K*_D_’s) from BLI were in the tens of nM regime, MST revealed no apparent binding under the same conditions (pH 7.5, 150 mM NaCl) (e.g. Fig S5a). Decreasing the salt to 20 mM NaCl, however, resulted in robust binding of a tandem N-peptide-GFP species, [N1-N2]-GFP, to Rubisco with a *K*_D_ of 75 nM on a stoichiometric binding site basis (i.e. one [N1-N2]-GFP binds to two of eight sites per Rubisco) (Fig. 2d).

MST indicated the N-peptide/Rubisco interaction is highly sensitive to salt concentration. Increasing NaCl from 20 mM to 60 mM showed a substantial increase in the *K*_D_ from 75 nM to 500 nM (Fig. 2e). Further increasing NaCl to 160 mM—near physiological ionic strength^32^ — weakened the binding beyond detection. Assuming a linear free energy relationship, we can estimate the binding energy for the individual N-peptide to Rubisco to be half of the ΔG^0^ for the [N1-N2]-GFP construct leading to *K*_D_ ∼ 250 μM at 20 mM NaCl (see SI, *MST fitting and analysis*). Indeed, MST of a single N-peptide-GFP, [N1]-GFP, showed no discernable binding over the same concentration range (Fig. S5b).

Taken together, these data present two puzzling observations. First, the individual N-peptide/Rubisco interaction alone appears too weak to drive carboxysome cargo encapsulation, particularly when approaching realistic intracellular ionic strength. Second, the relatively tight binding of Rubisco by a single N-peptide construct at 150 mM NaCl on BLI stands in apparent contradiction to the negative binding results obtained from MST under similar conditions. A mechanistic reconciliation of these issues is presented in the Discussion.

### Structural determination of the N-peptide/Rubisco complex

We next sought to obtain a structure of the N-peptide/Rubisco complex in order to locate the binding sites and to establish the nature of the specific molecular contacts. The NTD is largely disordered and its four N-peptide repeats could, in principle, adopt heterogeneous arrangements among the eight Rubisco binding sites. Furthermore, the binding of a single N-peptide is weak and salt sensitive. Disorder, structural heterogeneity, and partial occupancy therefore all pose significant challenges for co-crystallization. To circumvent these problems, we fused the N* consensus peptide to the C-terminus of the Rubisco large subunit (CbbL) via a short linker, -SS-, (Fig. 3a) to insure high local concentrations and saturation of all putative binding sites. This fusion protein was readily expressed, purified and was confirmed by size exclusion chromatography to be of the correct L_8_S_8_ oligomerization state (Fig. S6a). BLI measurements revealed no significant interaction of the Rubisco-N* fusion (prey) to surface N*-peptide (bait) suggesting that Rubisco-N* self-passivates its binding site (Fig. S6b,c).

**Figure 3.**
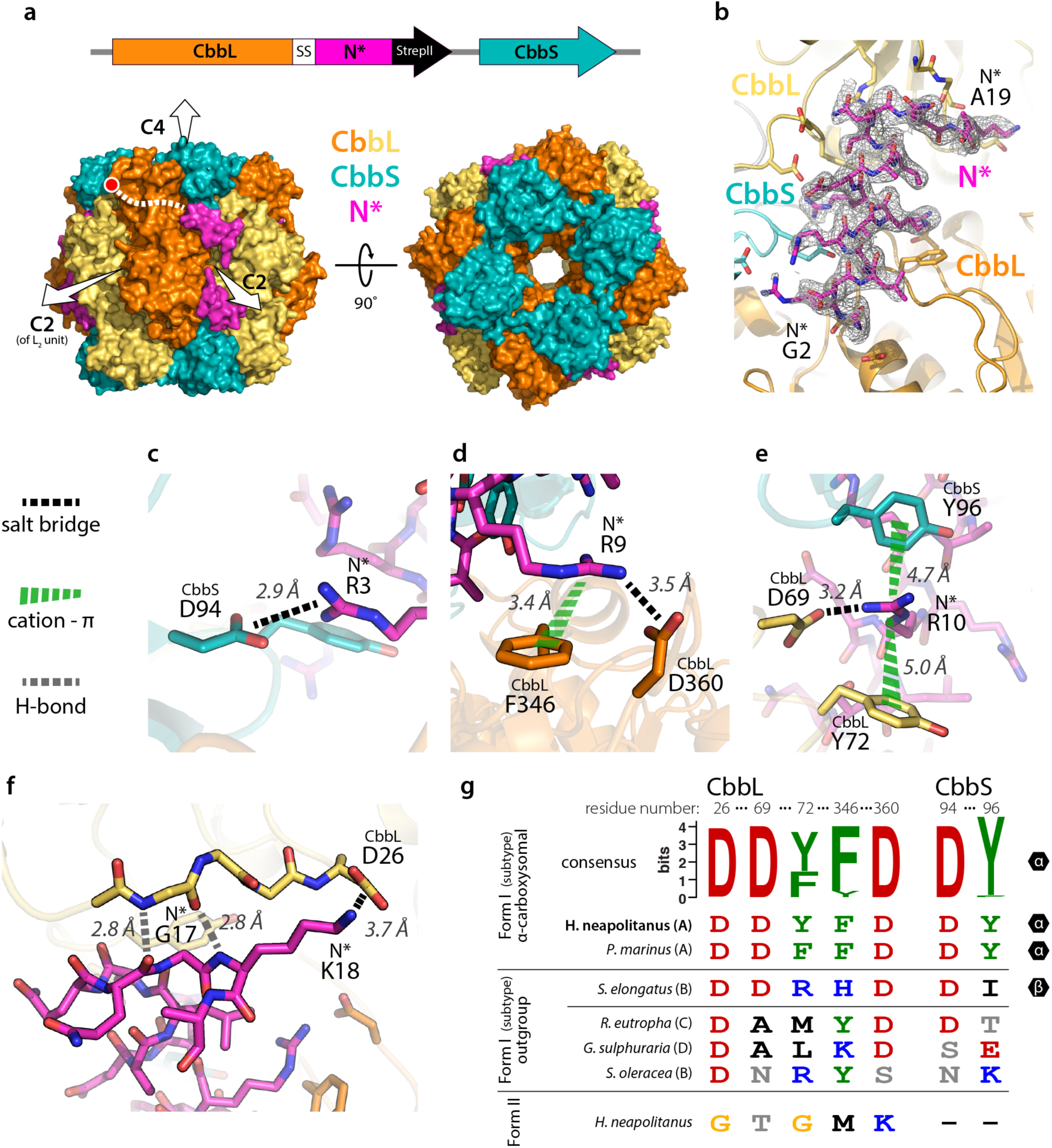
**a**, Schematic of the Rubisco-N* fusion construct and side and top views of a surface representation of the L_8_S_8_ biological assembly with bound N*-peptide. CbbL and CbbS are the large and small Rubisco subunits, respectively. The molecular symmetry axes are indicated by white arrows. The yellow and orange CbbLs are identical; the coloring is to highlight the L_2_ dimer units. The red dot is at the last structured residue of CbbL, while the dashed white line indicates the probable linkage to N*. **b**, Zoomed view of binding site with 2F_O_-F_C_ map at σ = 0.8 carved within 1.6 Å of N*. The first and last structured residues of N* are labeled. **c-f**, Molecular interactions of each of the five highly conserved residues of the N-peptide motif: R3, R9, R10, G17, and K18. Salt bridges, cation-π interactions, and select hydrogen bonds are specifically highlighted. The specific interactions were characterized with the PDBePISA33 and CaPTURE34 web servers. **g**, Rubisco sequence comparison at the N*-peptide interaction site. The Weblogo conservation sequence is from 231 α-carboxysomal Form IA Rubiscos. Two specific representatives, *H. neapolitanus* (used in this study) and *Prochlorococcus marinus MIT 9313*, are shown. Below are various outgroup Form I Rubiscos and the *H. neapolitanus* Form II Rubisco. Participation in carboxysomes (α or β) is indicated along the right of the table. Note that the residues are non-sequential and are numbered according to the *H. neapolitanus* sequence.

After screening and optimization of crystallization conditions, diffraction quality crystals were obtained (Table S1). X-ray diffraction data were collected and the structure was solved by molecular replacement using an existing model from Kerfeld and Yeates of *H. neapolitanus* Rubisco (PDB: 1SVD). The space group was C_2_ with four CbbL-N* and CbbS chains in the asymmetric unit. The Rubisco structure itself was essentially indistinguishable from wild-type with an average Cα RMSD of 0.27 Å. Clear unmodeled electron density was observed along the groove at the interface between two CbbL subunits (spanning separate L_2_ dimers) and a CbbS subunit (Fig. 3a). The N*-peptide was found to adopt a helical conformation and an all-atom model was manually built into the experimental density, which was sufficiently clear for unambiguous assignment of both the peptide direction and sequence registration. Following several rounds of refinement, the real-space cross-correlation for the modeled portion of N* (res. 2-19, Fig. 2c) was 90% or greater for each of the four N*-peptides in the asymmetric unit (Fig. 3b). All of the binding sites are occupied, indicating that the neighboring sites are not mutually occluding. Thus, it is likely that the L_8_S_8_ biological assembly possesses eight possible CsoS2 interaction sites.

The structure of the bound N*-peptide is largely α-helical, consistent with the secondary structure predictions and CD data (Fig. 1a,b). The last clearly structured residue of CbbL is at position 455, which is typical of structures of non-activated Form I Rubisco.^35^ The remainder of the CbbL C-terminus and the -SS-linker preceding N* are not observed in the electron density. Although lack of density complicates the assignment of N*/CbbL pairings, the structured portion of N* begins near CbbL helix 6 and the fusion thus likely originates from the C-terminus of this same subunit. This also agrees with previous structural models of other Rubiscos, in which the C-terminus extends over the so-called loop 6 in the same direction as the N* binding site (Fig 3a, dashed white).^35^ From there, the N* helix makes contacts with CbbS, spans the boundary to the neighboring L_2_ dimer, and finishes by breaking out of the helix at the N-terminal domain of the second CbbL. A noteworthy quality of the N*/Rubisco binding site is that, by contacting both CbbL and CbbS and bridging the L_2_ dimer interface, it exists only on the L_8_S_8_ Rubisco holoenzyme. This fact implies that only fully assembled Rubisco would be admitted into the carboxysome.

Each one of the highly conserved N* motif residues (Fig. 2c) is observed to make key binding contacts along the Rubisco interface. R3 is salt-bridged with CbbS D94 (Fig. 3c). R9 forms a salt-bridge with CbbL D360 and cation-π interaction with F346 (Fig. 3d). R10 has a salt-bridge to CbbL D69 and dual cation-π interactions with CbbL Y72 and CbbS Y96 (Fig. 3e). G17 appears to play a critical role in breaking the N* helix by facilitating backbone hydrogen bonds with CbbL and adopting glycine-specific Ψ-φ angles. Finally, K18 makes a salt bridge with CbbL D26 (Fig. 3f). All together the interactions are predominantly ionic and offer a structural explanation as to the energetic sensitivity to salt.

Amino acid residues involved in these electrostatic interactions are conserved for α-carboxysomal Form IA Rubisco. However, these residues were, in general, not conserved among an outgroup of various other Form I Rubiscos and the *H. neapolitanus* Form II Rubisco (Fig. 3g). To assay if these evolutionary observations are significant, two binding site mutants were made to test disruption of the binding interface. In one, each of the cation-π aromatics was mutated to alanine (CbbL Y72A, F346A; CbbS Y96A). In the other, a mutation was selected to resemble the β-carboxysomal Rubisco and to perturb the binding environment of N* R10 (CbbL Y72R). Neither mutant interacted with N* (Fig. S4).

### Structural comparison to CcmM/Rubisco

The general binding site of N*/Rubisco significantly overlaps with that of the recently determined CcmM/Rubisco interaction from the β-carboxysome, however, the specific molecular details are distinct.^17^ While CcmM binds with multiple regions across the SSUL domain,^17^ N* has a smaller footprint as a single α-helix (Fig. S10). In both cases, salt bridges—with the positive charge contributed by the scaffolding protein—are key parts of the interactions. A notable feature of the N*/Rubisco interaction, but absent in CcmM, are the prominent cation-π interactions.^34^ The complete conservation of the aromatics in the Rubisco binding site and the lack of binding when mutated to alanines suggest that the cation-π interactions indeed contribute meaningfully to the binding energy and specificity. Interestingly, cation-π contacts are a particularly common interaction modality among IDPs involved in protein liquid-liquid phase separation.^36–38^

### Hydrogen/deuterium exchange of carboxysomal versus purified Rubisco

To interrogate the CsoS2/Rubisco interaction in a native context, hydrogen/deuterium exchange (HDX) mass spectrometry experiments were performed in order to identify regions of Rubisco possessing differential protection when encapsulated within carboxysomes. HDX analysis of purified Rubisco versus carboxysomal Rubisco revealed a majority of peptides had nearly identical HDX rates. The most notable exception was CbbL 328-341 on helix 6 which experienced significantly greater protection inside carboxysomes (Fig. S8). This peptide, while not directly contacting N*, is connected through water-bridged hydrogen bond networks (Fig. S9). Although the NTD interaction does not apparently alter the crystal structure of Rubisco, it is possible that peptide binding may affect the dynamics of Rubisco structural elements.

### Effect of N-peptide multivalency on carboxysome formation

We set out to determine the importance of the number of N-peptide repeats on carboxysome assembly. *H. neapolitanus* CsoS2 contains four copies of the repeat but there is likely significant natural diversity. To this end, the consensus motif was used to quantify occurrences throughout the set of 231 CsoS2 genes.^39^ Every sequence contained at least two copies of the motif suggesting that a valency greater than one may be a general requirement for carboxysome assembly (Fig. 4b). Using a previously developed method whereby carboxysomes are produced heterologously in *E. coli* by expressing the known genes from a single plasmid (pHnCB10),^40^ we tested the effect of N-peptide repeat number on carboxysome formation. A series of pHnCB10 constructs were made possessing CsoS2 variants with a decreasing number of N-peptide repeats and tested for carboxysome expression. Only CsoS2 variants with two or more repeats were capable of forming carboxysomes (Fig. 4a and Fig. S11), consistent with the bioinformatic result.

**Figure 4.**
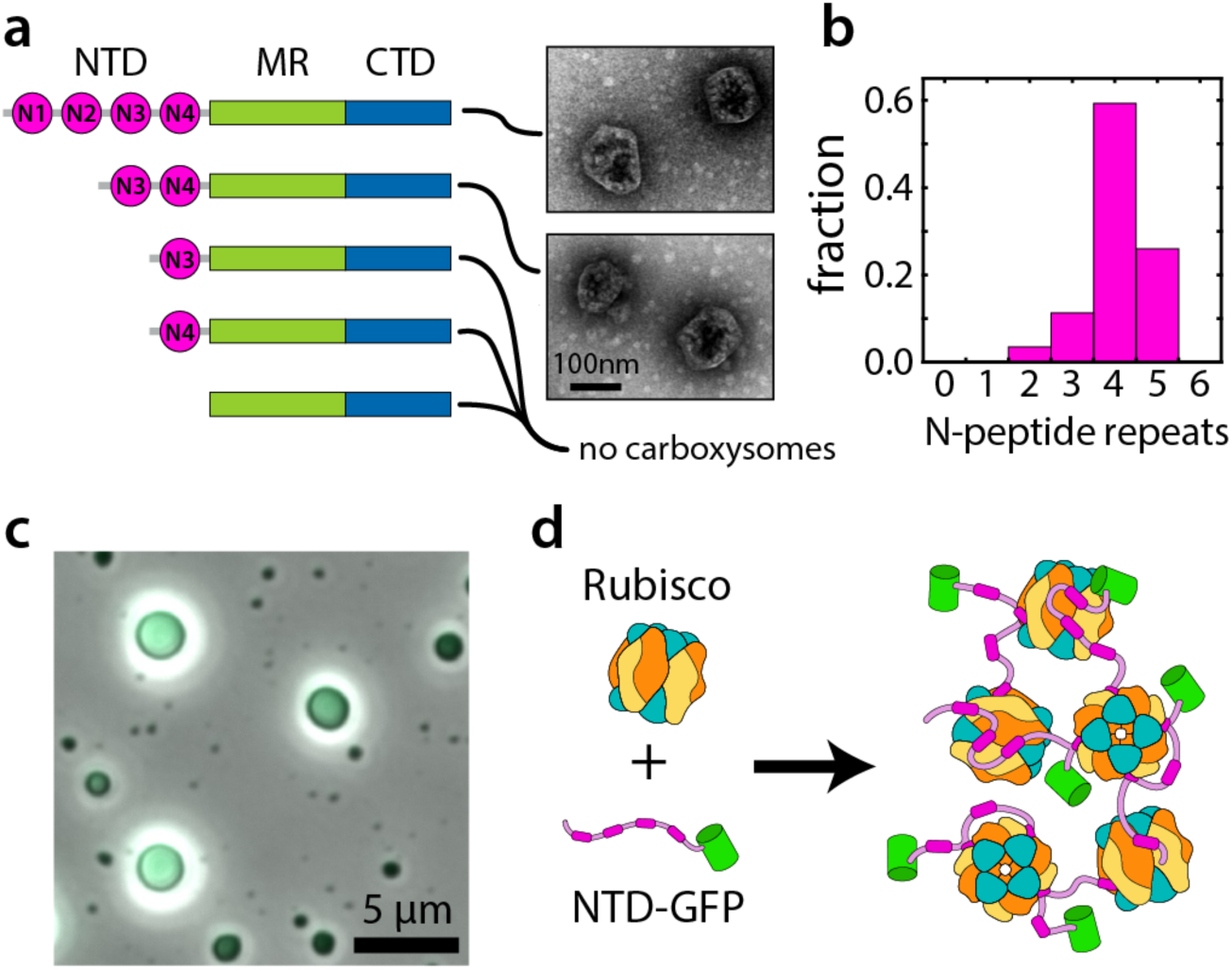
**a**, Truncated CsoS2 proteins with variable numbers of N-peptide repeats and TEM images of the resulting carboxysomes if any were formed. **b**, Histogram of N-peptide repeat numbers across 231 CsoS2 sequences. **c**, Merged GFP fluorescence and phase contrast images of protein liquid-liquid droplets formed from a solution of Rubisco and NTD-GFP. **d**, Microscopic model of the phase separated state. The branching of interactions due to the multivalency of both components provides the liquid cohesion while the relative weakness and exchangeability of the individual interactions confers fluidity.

### Phase separation of Rubisco and NTD

IDPs are highly represented in systems that undergo protein liquid-liquid phase separation. The propensity toward phase separation is promoted by weak individual interactions, often salt sensitive, and multivalent association either through well-defined binding sites or via less specific interactions related to the general amino acid composition.^41,42^ Phase separation has recently emerged as a common theme for the organization of Rubisco into CCM architectures. In the algal pyrenoid, Rubisco phase separates with EPYC1, a repetitive IDP.^43–45^ From β-carboxysomes the short form of the scaffold protein CcmM, M35,^46^ was shown to demix with Rubisco into protein liquid droplets.^17^ We hypothesized that CsoS2 and, in particular, the NTD may similarly demix with Rubisco. Indeed, when Rubisco and NTD-GFP are combined at 1.0 μM each at low salt (20 mM NaCl) the solution became turbid. Imaging by phase contrast and epifluorescence microscopy revealed that round green fluorescent droplets are formed (Fig. 4c) and are fully re-dissolved upon salt addition up to 150 mM NaCl. No droplets are observed with either individual component at the same concentrations.

## Discussion

We have characterized in molecular detail the binding interface of Rubisco and CsoS2 which facilitates α-carboxysome cargo encapsulation. CsoS2, as a large IDP, posed a significant challenge for structural determination. Through biophysical binding assays we narrowed down the interaction to a repeated motif within the CsoS2 NTD, fused this fragment directly to Rubisco, and obtained an x-ray crystal structure of the protein-peptide complex. We suggest that this workflow might be a valuable general strategy for determining structures of IDPs interacting with structured proteins since these interactions are often individually weak and transient.

Despite no apparent sequence similarity, the CsoS2/Rubisco binding bears striking parallels to the recently characterized CcmM/Rubisco interaction at the heart of β-carboxysome assembly.^17^ In both cases the scaffold protein binding element has multiple repeats interspersed by flexible linkers. The binding locations on Rubisco are very similar; both straddle an L_2_ dimer interface while also making critical contacts with a small subunit. This site is only present in the fully assembled L_8_S_8_ Rubisco holoenzyme so Rubisco assembly intermediates, namely L_2_ and (L_2_)_4_, would presumably not be encapsulated prematurely. Notwithstanding this global similarity, the specific structural details of the binding are distinct, making this an intriguing example of convergent evolution.

Another commonality between the α- and β-carboxysome scaffold/Rubisco systems is the propensity to undergo protein liquid-liquid phase separation. Phase separation is increasingly understood to play an organizational role in eukaryotes in the formation of membrane-less organelles.^47^ These structures and the droplets we observe (Fig. 4c), however, have at least a thousand-fold greater volume than carboxysomes. Furthermore, they are not enclosed within protein shells. Therefore, while suggestive of a dense liquid cargo phase, the role of demixing in the carboxysome assembly process remains unresolved.

The N-peptide/Rubisco interface is comprised chiefly of salt bridges and cation-π interactions. Consequently, the binding energy is highly sensitive to the solution ionic strength. Indeed, our solution phase binding measurements with MST indicate that the interaction dramatically weakens, with single site *K*_D_’s greater than 1 mM, at near-physiological ionic strength. Moreover, the phase separated droplets are fully dissolved under the same elevated salt concentrations. In apparent contradiction, however, the BLI measurements under the same conditions indicated strong binding (*K*_D_ ∼ 100 nM).

The essential difference is that BLI is a surface-based technique. Since the “prey” Rubsico has a site valency of eight, it could be simultaneously engaged by multiple “bait” N*-peptides in microscopically dense patches on the surface (see SI, *Comments on BLI*). This surface avidity effect enabled tight Rubisco binding even when the individual interactions were very weak. We propose that this artificial surface avidity represents a useful analogy to the early stages of carboxysome assembly. Several experiments have implicated CsoS2 association with the CsoS1 shell hexamer including native gel shifts^19^ and pulldown assays.^22^ Furthermore, the CsoS2 C-terminus was found at the shell^25^ and truncation of the CTD precludes carboxysome formation.^21^ Through the shell interaction, multiple CsoS2 molecules could be recruited to achieve high local concentration and then bind to Rubisco in a multivalent fashion with high affinity.

Our data have led us to the following speculative model of α-carboxysome assembly: At physiological ionic strength and the likely free concentrations of Rubisco and CsoS2 the interaction is insufficiently strong to drive significant association or demixing (Fig. 5a, point 1). However, in the presence of shell proteins, CsoS2 is gathered to high local concentration via interaction to the nascent shell surface and facilitates phase separation with Rubisco in the immediate vicinity of the shell (Fig. 5a, point 2). Eventually more shells with cargo droplets coalesce until the structure is fully enclosed.

**Figure 5.**
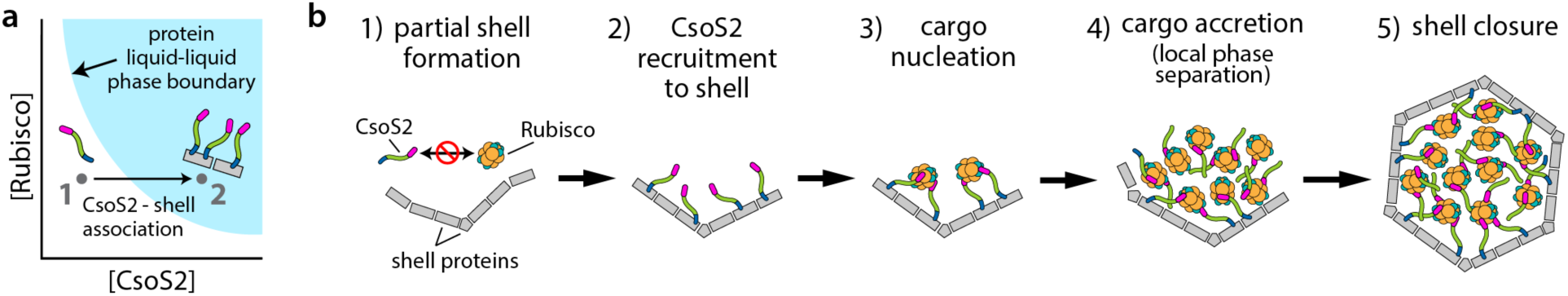
**a**, Model phase diagram of the hypothesized Rubisco/CsoS2 phase separation driven by the multivalent NTD interaction with Rubisco. The blue region represents the joint concentrations at which demixing occurs. At point 1 the cytosolic concentrations lie within the soluble region and both are fully dissolved. Through interactions with a nascent carboxysome shell, multiple CsoS2s are brought together, thus greatly increasing the concentration locally while the Rubisco concentration remains the same (point 2). This process locally exceeds the phase transition threshold and leads to local phase separation in the immediate vicinity of the shell. **b**, Model of α-carboxysome assembly in which the specific accumulation of cargo on the shell proceeds via the mechanism described in (**a**).

A full accounting of the interaction partners and the site binding energetics is alone insufficient to understand the carboxysome assembly process. Multivalency, surface avidity, protein liquid-liquid phase separation appear to play important roles but their relationships to the shell and the emergent size regularity remain unclear and warrant further investigation. Ultimately a detailed understanding of the principles of carboxysome assembly may be leveraged toward the design of synthetic microcompartments for biotechnological applications.

## Methods

### Protein expression and purification

All proteins used for biochemical assays contained a terminal affinity tag, either a hexahistidine tag or a Strep-tag II (see SI for complete sequences). Each construct was cloned via Golden Gate assembly ^48^ into a pET-14 based destination vector with ColE1 origin, T7 promoter, and carbenicillin resistance. These were transformed into *E. coli* BL21-AI expression cells. All Rubisco constructs were also co-transformed with pGro7 for expressing GroEL-GroES to facilitate proper protein folding. Cells were grown at 37°C to OD600 of 0.3-0.5 in 1 L of LB media before lowering the temperature to 18°C, inducing with 0.1% (w/v) L-arabinose, and growing overnight.

Cultures were harvested by centrifugation at 4,000 *g* and the pellets were frozen and stored at -80°C. The pellets were thawed on ice and resuspended with ∼25 mL of lysis buffer (50 mM Tris, 150 mM NaCl, pH 7.5) supplemented with 1 mM phenylmethanesulfonyl fluoride (PMSF), 0.1 mg/mL lysozyme, and 0.01 mg/mL DNaseI. The cells were lysed with three passes through an Avestin EmulsiFlex-C3 homogenizer and clarified by centrifugation at 12,000 *g* for 30 min. The clarified lysate was then incubated with the appropriate affinity resin for 30 min at 4°C with 2 mL of resin per 1 L of initial culture and transferred to a gravity column. His-tagged proteins were bound to HisPur Ni-NTA resin (Thermo), washed with lysis buffer with 30 mM imidazole, and eluted with lysis buffer with 300 mM imidazole. Strep-II-tagged proteins were bound to Strep-Tactin resin (EMD Millipore), washed with lysis buffer, and eluted with lysis buffer containing 2.5 mM desthiobiotin. All proteins were buffer exchanged to lysis buffer with 10DG Desalting Columns (Bio-Rad). For storage, proteins were made to 10% (w/v) glycerol, flash frozen in liquid nitrogen, and stored at -80°C.

Protein purity was assessed by SDS-PAGE gel analysis. In general all protein was >90% the desired product. Size exclusion chromatography was performed analytically to confirm purity and aggregation state and, if needed, as a final preparative step.

### Bio-layer interferometry

Protein-protein interactions were measured using bio-layer interferometry (BLI) with an Octet RED384 (Forte Bio). The “bait” protein was immobilized on Ni-NTA Dip and Read Biosensors via a terminal His-tag. Typical “bait” concentrations for the sensor loading were 10 μg/mL. The soluble “prey” protein concentrations were varied in the nanomolar to micromolar range. The buffer used for all loading, association/dissociation, and wash steps was 50 mM Tris, 150 mM NaCl, 0.01% (w/v) Triton X-100, pH 7.5. Sensor regeneration of the Ni-NTA was done with 50 mM Tris, 150 mM NaCl, 0.05% (w/v) SDS, 300 mM imidazole, pH 7.5. The typical experimental binding sequence used was: load “bait”, buffer wash, “prey” association, “prey” dissociation in buffer, sensor regeneration, buffer wash. For the experiments testing the binding activity of specific peptides (Fig. 2b and Fig. S2), “bait” proteins were designed with a 40 amino acid proline rich region between the His-tag and the peptide (see SI, *Protein Sequences*). This insertion is expected to adopt an extended polyproline II helix conformation ∼10 nm in length ^49^ and was included to limit possible surface occlusion.

### Microscale thermophoresis

Solution protein-protein binding was monitored by microscale thermophoresis (MST) with a Monolith NT.115 (Nanotemper). The target proteins were portions of the CsoS2 NTD fused to Superfolder GFP and used at a concentration of 50 nM. Unlabeled Rubisco was used as the ligand with concentrations varied in two-fold increments from 10 μM (as L_8_S_8_) down to 0.3 nM. Experiments were carried out in buffer with 6.7 mM Tris, 0.01% Triton X-100, pH 7.5 and either 20, 60, or 160 mM NaCl. The samples were loaded into MST Premium Coated Capillaries (Nanotemper) and analyzed using 20% blue LED power for fluorescence excitation and Medium infrared laser power for the thermophoresis. Data fitting and bootstrap error estimation was performed using custom scripts in MATLAB (MathWorks).

### Crystallization, x-ray diffraction, and refinement

Initial screening of crystallization conditions for CbbL-N*, CbbS was done using the Hampton Crystal Screen (HR2-110) with protein at 15 mg/mL combined 1:1 with the screen mother liquors. Due to the hypothesized ionic nature of the interaction, screen conditions having lower salt concentrations were prioritized in the follow-up optimization. Ultimately the best crystals were obtained from a mother liquor of 0.2M MgCl_2_ • 6H_2_O, 0.1M HEPES, 30% (v/v) PEG-400. Protein at 15 mg/mL diluted 1:2 with mother liquor was allowed to equilibrate for one day by hanging drop vapor diffusion whereupon it was microseeded with pulverized crystals from more concentrated conditions delivered with a cat whisker.

Crystals were looped and directly frozen on the beamline under a 100K nitrogen jet without additional cryoprotectant. X-ray diffraction was collected with wavelength 1.11 Å on a Pilatus3 S 6M (Dectris) detector with a 50μm beam pinhole at the Advanced Light Source, BL 8.3.1, Berkeley, CA.

The data were indexed and integrated with XDS^50^ and scaled and merged with AIMLESS.^51,52^ Molecular replacement was carried out in Phenix using the existing wild-type *H. neapolitanus* Rubisco structure (PDB ID: 1SVD) as the search model.^53,54^ Cycles of automatic refinement were performed with Phenix while Coot was used for manual model building.^55^ The final refined structure backbone conformations were 96.0% Ramachandran favored, 3.8% allowed, and 0.2% outliers.

### Carboxysome construct generation and purification

Heterologous expression of carboxysomes in *E. coli* was performed following the methods of Bonacci et al. using the plasmid pHnCB10 which contains genes encoding all ten of the proteins known to participate in carboxysome formation.^40^ Golden Gate assembly was used to make the truncations of the CsoS2 NTD shown in Fig. 4a.

Carboxysomes were purified as previously described.^21^ Briefly, the cells were harvested, resuspended in 25mL TEMB buffer (10 mM Tris, 10 mM MgCl_2_, 1 mM EDTA, and 20 mM NaHCO_3_, pH 8.4), lysed with a homogenizer, and the lysate clarified by centrifugation at 12,000 *g* for 30 min. The supernatant was further centrifuged at 40,000 *g* for 30min to pellet the carboxysomes. The carboxysome pellet was resuspended in 1x Cellytic B (Sigma-Aldrich) in order to solubilize any residual membrane fragments. The solution was spun a second time at 40,000 *g* for 30 min to pellet the carboxysomes again. The pellet was resuspended with 3mL of TEMB, clarified with a 5min spin at 3,000 *g*, and loaded on top of a 25-mL sucrose step gradient (10, 20, 30, 40, and 50% w/v sucrose). This was ultracentrifuged at 105,000 *g* for 30 min. The solution was fractionated and analyzed by SDS-PAGE. Those fractions containing the expected set of carboxysomal proteins (and which also demonstrated visible Tyndall scattering) were pooled, pelleted by centrifugation for 90min at 105,000 *g*, resuspended in 1mL of TEMB, and stored at 4°C.

### Negative stain TEM

Purified carboxysomes were visualized by negative stain transmission electron microscopy. Formvar/carbon coated copper grids were prepared by glow discharge prior to sample application. The grids were washed with deionized water several times before staining with 2% (w/v) uranyl acetate. Imaging was performed on a JEOL 1200 EX transmission electron microscope.

### Hydrogen/deuterium exchange mass spectrometry

Peptide mass fingerprinting from purified Rubisco and carboxysomes was performed using on-column pepsin digestion, followed by reversed-phase HPLC, and tandem mass spectrometry on a Thermo Scientific LTQ Orbitrap Discovery.^56,57^ For hydrogen exchange, the samples were diluted 1:10 in D_2_O buffer (50 mM Tris, 150 mM NaCl, pD 7.5) and then aliquots removed and quenched in 500 mM glycine, 2 M guanidinium hydrochloride (GdnHCl), pH 2.0 buffer at log-spaced time intervals from 20 seconds to 48 hours. Samples were immediately frozen in liquid nitrogen upon addition of quenching solution. Deuterated control samples were prepared by 1:10 dilution in D_2_O, 50 mM Tris, 150 mM NaCl, 6 M GdnHCl, pD 7.5 and quenching with 500 mM glycine, pH 2.0. Samples were thawed, digested on-column as before, and analyzed by LCMS. Data analysis was performed with HDExaminer (Sierra Analytics).

### CD spectroscopy

Purified protein was first exchanged into CD buffer (20 mM sodium phosphate and 20 mM sodium sulfate, pH 7.4) to minimize the background absorbance. From this solution, 300 µL was transferred to a 1-mm quartz cell. The sample containing only CD buffer was included as a negative control. Data were collected on a J-815 circular dichroism spectrometer (JASCO). Spectra were collected from 190 to 260 nm in 0.5 nm steps with the scanning speed of 20 nm/min and signal averaging for 1 s for each step. Each sample was measured 3 times and the spectra were averaged. Protein concentrations were determined using 280 nm absorbance and extinction coefficients calculated using ProtParam.

### Bioinformatics

The CsoS2 secondary structure predictions were made using JPred.^27^ The disorder score was calculated with PONDR-FIT.^26^

The candidate α-carboxysome-associated CsoS2 sequences were selected from the Integrated Microbial Genomes (IMG) database by searching for the CsoS2 PFAM (PF12288) within 100kb of loci containing the Rubisco large and small subunits (PF00016 and PF00101), α-carboxysomal carbonic anhydrase (PF08936), and bacterial microcompartment shell proteins (PF00936). These sequences (n=231) were aligned with ClustalOmega,^58^ truncated to include only the NTD (i.e. all sequence before the first MR repeat), and analyzed with MEME ^31^ to find repeated sequence motifs (Fig. 2c). The Motif Alignment and Search Tool (MAST) ^39^ was used to locate and count all occurrences of the motif within the full CsoS2 sequences (Fig. 4b).

## Acknowledgements

We thank Cecilia Blikstad for helpful comments on the manuscript. We also thank Peter Huang for his help with the BLI instrumentation and Cheryl Kerfeld for advice on Rubisco crystallization. Yinon Bar-On assisted us in gathering the CsoS2 sequences. We acknowledge the staff at the UC Berkeley Electron Microscope Laboratory for training and assistance with TEM. George Meigs and James Holton assisted with the x-ray diffraction and we gratefully acknowledge their input. Whiskers for crystal microseeding were kindly gifted by S.T. Kuhl. Beamline 8.3.1 at the Advanced Light Source is operated by the University of California Office of the President, Multicampus Research Programs and Initiatives grant MR-15-328599, the National Institutes of Health (R01 GM124149 and P30 GM124169), Plexxikon Inc. and the Integrated Diffraction Analysis Technologies program of the US Department of Energy Office of Biological and Environmental Research. The work was supported by grants from the U.S. Department of Energy (DE-SC00016240) and the National Institute of General Medical Sciences (R01GM129241) to D.F.S. and a grant from the National Institute of General Medical Sciences (R01GM050945) to S.M.

## References

1. Raven, J. A., Cockell, C. S. & De La Rocha, C. L. The evolution of inorganic carbon concentrating mechanisms in photosynthesis. Philos. Trans. R. Soc. Lond. B. Biol. Sci 363, 2641–2650 (2008).

2. Mangan, N. M., Flamholz, A., Hood, R. D., Milo, R. & Savage, D. F. pH determines the energetic efficiency of the cyanobacterial CO2 concentrating mechanism. Proc Natl Acad Sci USA 113, E5354–62 (2016).

3. Espie, G. S. & Kimber, M. S. Carboxysomes: cyanobacterial RubisCO comes in small packages. Photosyn. Res. 109, 7–20 (2011).

4. Rae, B. D., Long, B. M., Badger, M. R. & Price, G. D. Functions, compositions, and evolution of the two types of carboxysomes: polyhedral microcompartments that facilitate CO2 fixation in cyanobacteria and some proteobacteria. Microbiol. Mol. Biol. Rev. 77, 357–379 (2013).

5. Heinhorst, S., Cannon, G. C. & Shively, J. M. in Complex intracellular structures in prokaryotes (ed. Shively, J. M.) 2, 141–165 (Springer Berlin Heidelberg, 2006).

6. Kerfeld, C. A. & Melnicki, M. R. Assembly, function and evolution of cyanobacterial carboxysomes. Curr. Opin. Plant Biol. 31, 66–75 (2016).

7. Tanaka, S. et al. Atomic-level models of the bacterial carboxysome shell. Science 319, 1083–1086 (2008).

8. Schmid, M. F. et al. Structure of Halothiobacillus neapolitanus carboxysomes by cryoelectron tomography. J. Mol. Biol. 364, 526–535 (2006).

9. Iancu, C. V. et al. The structure of isolated Synechococcus strain WH8102 carboxysomes as revealed by electron cryotomography. J. Mol. Biol. 372, 764–773 (2007).

10. Shih, P. M. et al. Biochemical characterization of predicted Precambrian RuBisCO. Nat. Commun. 7, 10382 (2016).

11. Whitehead, L., Long, B. M., Price, G. D. & Badger, M. R. Comparing the in vivo function of α-carboxysomes and β-carboxysomes in two model cyanobacteria. Plant Physiol. 165, 398–411 (2014).

12. Shively, J. M., Ball, F., Brown, D. H. & Saunders, R. E. Functional organelles in prokaryotes: polyhedral inclusions (carboxysomes) of Thiobacillus neapolitanus. Science 182, 584–586 (1973).

13. Cameron, J. C., Wilson, S. C., Bernstein, S. L. & Kerfeld, C. A. Biogenesis of a bacterial organelle: the carboxysome assembly pathway. Cel. 155, 1131–1140 (2013).

14. Kinney, J. N., Salmeen, A., Cai, F. & Kerfeld, C. A. Elucidating essential role of conserved carboxysomal protein CcmN reveals common feature of bacterial microcompartment assembly. J. Biol. Chem. 287, 17729–17736 (2012).

15. Long, B. M., Badger, M. R., Whitney, S. M. & Price, G. D. Analysis of carboxysomes from Synechococcus PCC7942 reveals multiple Rubisco complexes with carboxysomal proteins CcmM and CcaA. J. Biol. Chem. 282, 29323–29335 (2007).

16. Ryan, P. et al. The small RbcS-like domains of the β-carboxysome structural protein CcmM bind RubisCO at a site distinct from that binding the RbcS subunit. J. Biol. Chem. 294, 2593–2603 (2019).

17. Wang, H. et al. Rubisco condensate formation by CcmM in β-carboxysome biogenesis. Nature 566, 131–135 (2019).

18. Long, B. M., Rae, B. D., Badger, M. R. & Price, G. D. Over-expression of the β-carboxysomal CcmM protein in Synechococcus PCC7942 reveals a tight co-regulation of carboxysomal carbonic anhydrase (CcaA) and M58 content. Photosyn. Res. 109, 33–45 (2011).

19. Cai, F. et al. Advances in understanding carboxysome assembly in prochlorococcus and synechococcus implicate csos2 as a critical component. Life (Basel. 5, 1141–1171 (2015).

20. Cannon, G. C. et al. Organization of carboxysome genes in the thiobacilli. Curr. Microbiol. 46, 115–119 (2003).

21. Chaijarasphong, T. et al. Programmed Ribosomal Frameshifting Mediates Expression of the α-Carboxysome. J. Mol. Biol. 428, 153–164 (2016).

22. Williams, E. B. Identification and Characterization of Protein Interactions in the Carboxysome of Halothiobacillus neapolitanus. (2006).

23. Liu, Y. et al. Deciphering molecular details in the assembly of alpha-type carboxysome. Sci. Rep. 8, 15062 (2018).

24. Gonzales, A. D. et al. Proteomic analysis of the CO2-concentrating mechanism in the open-ocean cyanobacteriumSynechococcus WH8102. Can. J. Bot. 83, 735–745 (2005).

25. Baker, S. H. et al. The correlation of the gene csoS2 of the carboxysome operon with two polypeptides of the carboxysome in thiobacillus neapolitanus. Arch. Microbiol. 172, 233–239 (1999).

26. Xue, B., Dunbrack, R. L., Williams, R. W., Dunker, A. K. & Uversky, V. N. PONDR-FIT: a meta-predictor of intrinsically disordered amino acids. Biochim. Biophys. Acta 1804, 996–1010 (2010).

27. Drozdetskiy, A., Cole, C., Procter, J. & Barton, G. J. JPred4: a protein secondary structure prediction server. Nucleic Acids Res. 43, W389–94 (2015).

28. Abdiche, Y., Malashock, D., Pinkerton, A. & Pons, J. Determining kinetics and affinities of protein interactions using a parallel real-time label-free biosensor, the Octet. Anal. Biochem. 377, 209–217 (2008).

29. van der Lee, R. et al. Classification of intrinsically disordered regions and proteins. Chem. Rev. 114, 6589–6631 (2014).

30. Davey, N. E. et al. Attributes of short linear motifs. Mol. Biosyst. 8, 268–281 (2012).

31. Bailey, T. L. & Elkan, C. Fitting a mixture model by expectation maximization to discover motifs in biopolymers. Proc. Int. Conf. Intell. Syst. Mol. Biol. 2, 28–36 (1994).

32. Alberty, R. A. Thermodynamics of biochemical reactions. (John Wiley & Sons, Inc., 2003). doi:10.1002/0471332607

33. Krissinel, E. & Henrick, K. Inference of macromolecular assemblies from crystalline state. J. Mol. Biol. 372, 774–797 (2007).

34. Gallivan, J. P. & Dougherty, D. A. Cation-pi interactions in structural biology. Proc Natl Acad Sci USA 96, 9459–9464 (1999).

35. Schneider, G., Lindqvist, Y. & Brändén, C. I. RUBISCO: structure and mechanism. Annu. Rev. Biophys. Biomol. Struct. 21, 119–143 (1992).

36. Brangwynne, C. P., Tompa, P. & Pappu, R. V. Polymer physics of intracellular phase transitions. Nat. Phys. 11, 899–904 (2015).

37. Nott, T. J. et al. Phase transition of a disordered nuage protein generates environmentally responsive membraneless organelles. Mol. Cell 57, 936–947 (2015).

38. Qamar, S. et al. FUS Phase Separation Is Modulated by a Molecular Chaperone and Methylation of Arginine Cation-π Interactions. Cel. 173, 720-734.e15 (2018).

39. Bailey, T. L. & Gribskov, M. Combining evidence using p-values: application to sequence homology searches. Bioinformatics 14, 48–54 (1998).

40. Bonacci, W. et al. Modularity of a carbon-fixing protein organelle. Proc Natl Acad Sci USA 109, 478–483 (2012).

41. Li, P. et al. Phase transitions in the assembly of multivalent signalling proteins. Nature 483, 336–340 (2012).

42. Boeynaems, S. et al. Protein phase separation: A new phase in cell biology. Trends Cell Biol. 28, 420–435 (2018).

43. Mackinder, L. C. M. et al. A repeat protein links Rubisco to form the eukaryotic carbon-concentrating organelle. Proc Natl Acad Sci USA 113, 5958–5963 (2016).

44. Wunder, T., Cheng, S. L. H., Lai, S.-K., Li, H.-Y. & Mueller-Cajar, O. The phase separation underlying the pyrenoid-based microalgal Rubisco supercharger. Nat. Commun. 9, 5076 (2018).

45. Freeman Rosenzweig, E. S. et al. The Eukaryotic CO2-Concentrating Organelle Is Liquid-like and Exhibits Dynamic Reorganization. Cel. 171, 148-162.e19 (2017).

46. Long, B. M., Tucker, L., Badger, M. R. & Price, G. D. Functional cyanobacterial beta-carboxysomes have an absolute requirement for both long and short forms of the CcmM protein. Plant Physiol. 153, 285–293 (2010).

47. Hyman, A. A., Weber, C. A. & Jülicher, F. Liquid-liquid phase separation in biology. Annu. Rev. Cell Dev. Biol. 30, 39–58 (2014).

48. Engler, C., Kandzia, R. & Marillonnet, S. A one pot, one step, precision cloning method with high throughput capability. PLoS ONE 3, e3647 (2008).

49. Schuler, B., Lipman, E. A., Steinbach, P. J., Kumke, M. & Eaton, W. A. Polyproline and the “spectroscopic ruler” revisited with single-molecule fluorescence. Proc Natl Acad Sci USA 102, 2754–2759 (2005).

50. Kabsch, W. Integration, scaling, space-group assignment and post-refinement. Acta Crystallogr. D Biol. Crystallogr. 66, 133–144 (2010).

51. Collaborative Computational Project, Number 4. The CCP4 suite: programs for protein crystallography. Acta Crystallogr. D Biol. Crystallogr. 50, 760–763 (1994).

52. Evans, P. R. & Murshudov, G. N. How good are my data and what is the resolution? Acta Crystallogr. D Biol. Crystallogr. 69, 1204–1214 (2013).

53. McCoy, A. J. et al. Phaser crystallographic software. J. Appl. Crystallogr. 40, 658–674 (2007).

54. Adams, P. D. et al. PHENIX: a comprehensive Python-based system for macromolecular structure solution. Acta Crystallogr. D Biol. Crystallogr. 66, 213–221 (2010).

55. Emsley, P. & Cowtan, K. Coot: model-building tools for molecular graphics. Acta Crystallogr. D Biol. Crystallogr. 60, 2126–2132 (2004).

56. Lim, S. A., Bolin, E. R. & Marqusee, S. Tracing a protein’s folding pathway over evolutionary time using ancestral sequence reconstruction and hydrogen exchange. elife 7, (2018).

57. Samelson, A. J. et al. Kinetic and structural comparison of a protein’s cotranslational folding and refolding pathways. Sci. Adv. 4, eaas9098 (2018).

58. Sievers, F. & Higgins, D. G. Clustal Omega for making accurate alignments of many protein sequences. Protein Sci. 27, 135–145 (2018).

