## Supplemental Information for "α-carboxysome formation is mediated by the multivalent and disordered protein CsoS2"

### Supplementary Information:

*Comments on bio-layer interferometry (BLI)* 2

*BLI of select CsoS2 peptides with Rubisco* 4

*BLI of mutant NTD* 5

*Rubisco binding site mutants* 6

*Microscale thermophoresis (MST) fitting and analysis* 7

*Rubisco-N* fusion characterization* 9

*X-ray crystallography refinement statistics* 10

*Enrichment of binding motif from peptide array data* 11

*Hydrogen / deuterium exchange of Rubisco inside and outside carboxysomes* 13

*Structural comparison to CcmM/Rubisco interaction* 14

*CsoS2 NTD truncations and carboxysome formation* 18

*Protein sequences* 19

*Plasmids used* 25

*Comments on bio-layer interferometry (BLI)*

Bio-layer interferometry (BLI) played a critical role in this study as a robust, medium-throughput way to narrow down the CsoS2/Rubisco binding activity to the N-peptide and as a qualitative test of the effects of binding site mutations on the activity. In principle this method can yield specific information on binding energies. However, in this case, the high valency of the interaction (8 for Rubisco and 4 for CsoS2) combined with surface avidity effects make this kind of energetic determination infeasible.

For any surface-based binding measurement, false positives due to non-specific binding are a concern. Our confidence in the qualitative binding results is born of two observations: one, there is very little signal accumulation on unloaded biosensors (e.g. Fig. S2, black trace) and, two, minor targeted modifications to the bait or the prey could entirely abolish binding (e.g. randomizing the N*-peptide sequence eliminated all Rubisco binding; see Fig. 2b).

The BLI data for every bait construct with binding activity towards Rubisco (i.e. full length CsoS2, NTD, N*-polyPro) had clear signatures of surface avidity. This situation is essentially unavoidable because Rubisco has a valency of 8 and the BLI biosensors require high surface densities relative to Rubisco's size. In the case of CsoS2 and NTD, they both contain four N-peptides and the avidity effects are particularly acute as evidenced by the heterogeneous dissociation kinetics (Fig. 2a) which likely arise from density variations of bait and result in microscopic surface sites of differing affinities. N*-polyPro, with just a single N-peptide, demonstrates simpler fully reversible binding but is nevertheless still poorly modeled as a 1:1 binding interaction (Fig. S1). The implied dissociation constant is around 100 nM but it is not clear, for example, how many individual interactions are involved. In any event, we make no attempt to quantify the binding energetics from these results but rather took them as qualitative indications of binding.

The data in Fig. S1 was globally fit with the following piece-function:

Eq. S1
$$0 < t \leq t_{dissoc}, \quad S(t, c_{Ru}) = S_{sat} * \left( \frac{1 - e^{-(k_a c_{Ru} + k_d)\, t}}{1 + \frac{k_d}{k_a c_{Ru}}} \right)$$

$$t_{dissoc} < t, \qquad S(t, c_{Ru}) = S(t_{dissoc}, c_{Ru}) * e^{-k_d (t - t_{dissoc})}$$

where $t$ is time, $t_{dissoc}$ is the time of biosensor transfer to dissociation buffer, $c_{Ru}$ is the Rubisco concentration, $k_a$ is the bimolecular kinetic association rate constant, $k_d$ is the unimolecular dissociation rate constant, and $S_{sat}$ is the BLI signal at saturation.

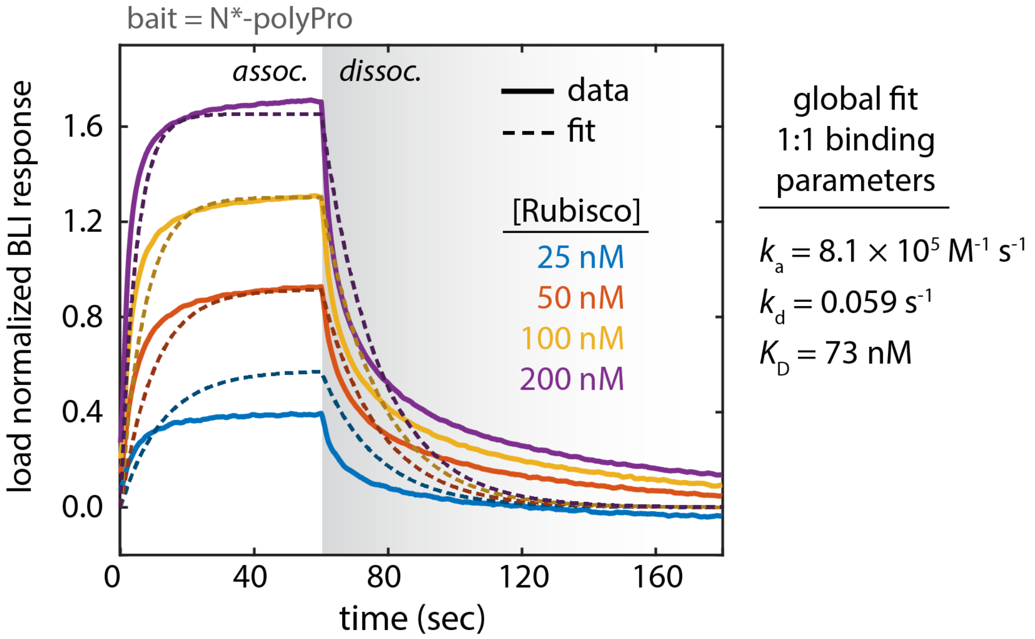

**Figure S1**

BLI response curves for Rubisco binding to N*-polyPro. The 1:1 binding model global fit to Eq. S1 is shown. Due to the high valency of Rubisco, this model only applies in the limit of very low surface density of the monovalent bait (N*-polyPro). The binding profiles exhibit deviations from this idealization which are consistent with significant surface avidity effects. It therefore does not lend itself to simple deconvolution of the energetics.

*BLI of select CsoS2 peptides with Rubisco*

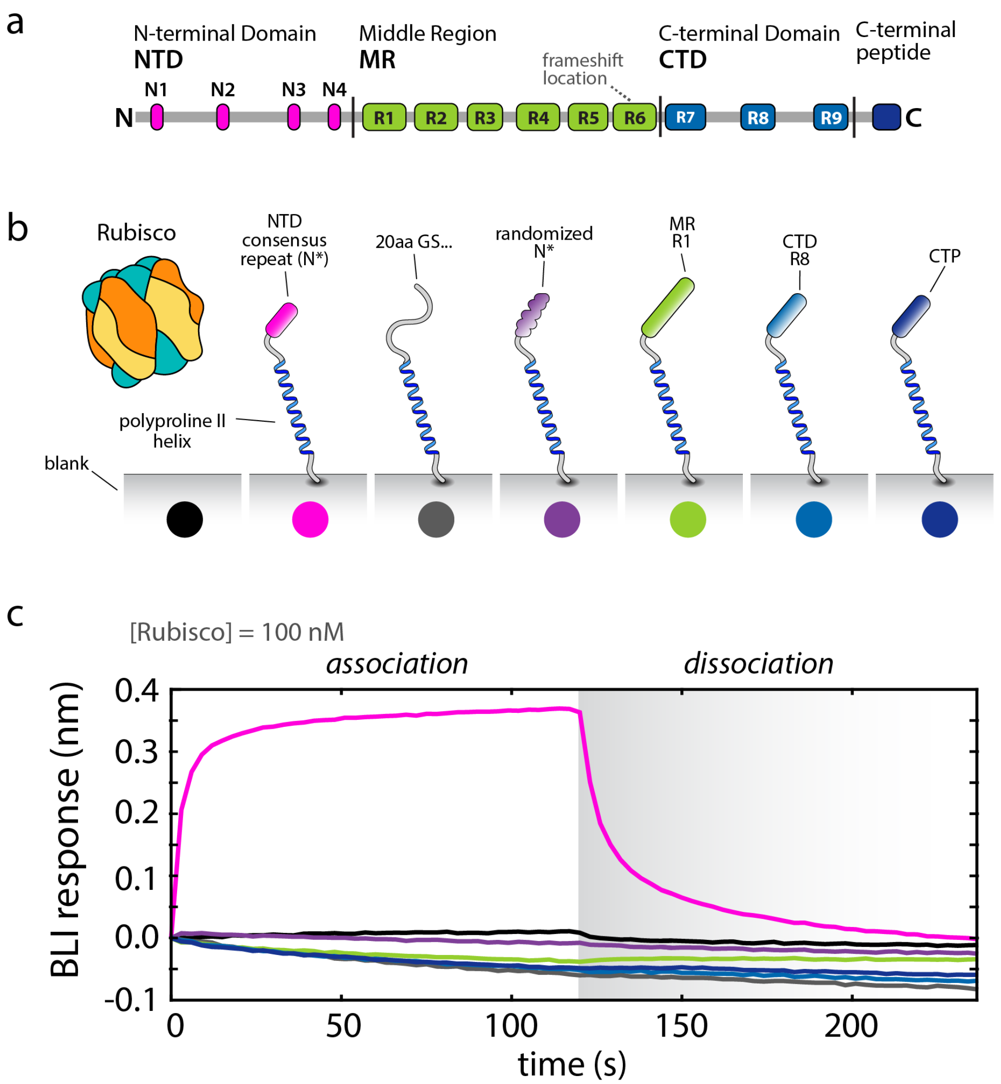

**Figure S2**

**a**, Primary sequence of CsoS2 with each of the repeated or conserved elements. **b**, Schematic representation of a set of BLI experiments testing the specificity of the Rubisco - CsoS2 interaction. Each of a series of CsoS2 elements and control sequences was fused to polyproline II helices which were surface immobilized to a Ni-NTA functionalized biosensor surface via an N-terminal hexahistidine tag. **c**, BLI traces of the constructs from (**b**) when incubated with 100 nM Rubisco. The trace colors match the dots in (**b**). Only the N*-peptide demonstrates any specific binding activity.

*BLI of mutant NTD*

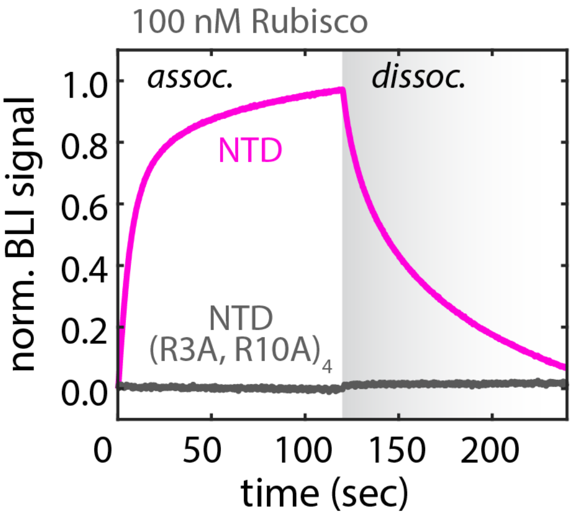

**Figure S3**

BLI response towards 100 nM Rubisco with bait of either the NTD or the NTD with R3A, R10A mutations made within all four of the N-peptide repeats. Removing those conserved arginines entirely eliminates the binding.

*Rubisco binding site mutants*

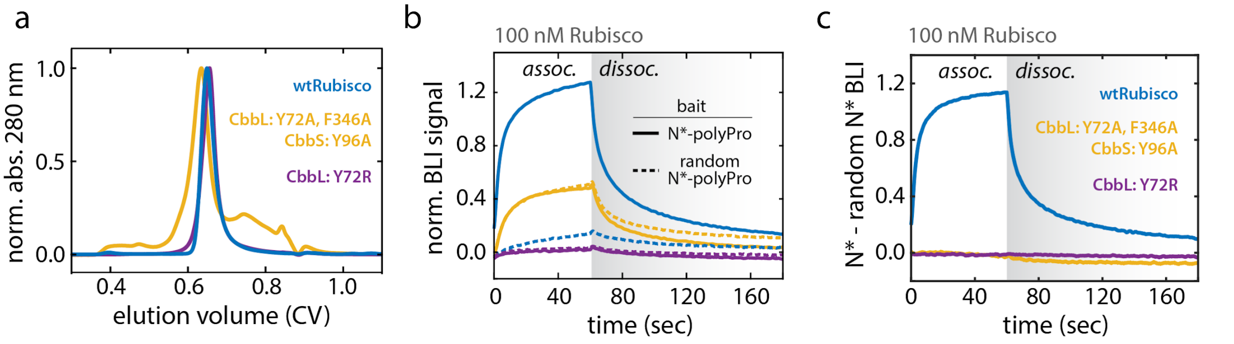

**Figure S4**

**a**, Size exclusion chromatograms of wild-type *H. neapolitanus* Rubisco (wtRubisco), a mutant with all cation-π aromatics mutated to alanines (CbbL: Y72A, F346A; CbbS: Y96A), and a salt bridge disrupting mutation (CbbL: Y72A). All species eluted at a volume consistent with the $L_8S_8$ structure. **b**, Each Rubisco species was tested for binding activity by BLI to N*-polyPro (solid lines) and the randomized N*-polyPro negative control (dashed lines). Only the wild-type Rubisco had specific binding activity to N*-polyPro over the randomized N*-peptide control. The aromatic removal mutant (yellow) had some non-specific binding to both baits but showed no preference for the real N*-peptide sequence. **c**, Differential BLI binding signal of each Rubisco species to N*-polyPro relative to random N*-polyPro. Both Rubisco binding site mutants clearly possess no specific association.

*Microscale thermophoresis (MST) fitting and analysis*

MST data were, in general, collected for a series of 16 ligand concentrations as serial 2-fold dilutions. The target concentrations were 50 nM of the GFP fusion species. The isotherms were fit to a 1:1 binding model according to the law of mass action,

Eq. S2

$$S(c_L) = S_{unbound} + (S_{bound} - S_{unbound}) * \frac{c_L - c_T + K_D - \sqrt{(c_L + c_T + K_D)^2 - 4\, c_L\, c_T}}{2\, c_T}$$

where $S$ is the observed MST response, $S_{\text{unbound}}$ and $S_{\text{bound}}$ are the MST responses of the free and saturated target respectively, $c_{\text{L}}$ is the ligand concentration (varied over 16 concentrations), $c_{\text{T}}$ is the target concentration (constant for each experiment), and $K_{\text{D}}$ is the dissociation constant.

The ligand concentration is taken as the effective total concentration of binding sites available to the target. For example, [N1-N2]-GFP can engage two of the eight sites on Rubisco, therefore, the ligand concentration is four times the Rubisco holoenzyme ($L_8S_8$) concentration. Implicit in this simple treatment are a number of assumptions. One, the targets bind with all N-peptides (e.g. 2 for [N1-N2]-GFP). Two, all possible binding configurations (that is, the microscopic arrangements of the N-peptide binding locations allowable by the linking region) have equivalent binding energies. And three, the thermophoretic response of the bound species will be the same regardless of the number of targets bound to that particular Rubisco. Deviations from these assumptions are expected to be small and do not justify the inclusion of additional fit parameters.

Where applicable the fitting procedure was conducted by taking the median fit parameters from bootstrap sampling. Specifically, a subset of the data was randomly selected from among the experimental replicates. This subset was fit to Eq. S2 by least-squares fitting and the parameters recorded. The process was repeated 10,000 times. The mean and confidence intervals for $K_{\text{D}}$ were determined from the resulting distribution.

The single site (i.e. one N-peptide to one Rubisco site) binding energy proved to be too weak to accurately determine under our experimental conditions (see Fig. S5b). Consequently, we made rough estimates of the single site binding constants by assuming a linear free energy relationship, that is, the free energy of binding for a single site was taken as half that of the bivalent species ([N1-N2]-GFP; see Fig. 2e dashed lines). This should be construed as an upper limit since multivalent ligands are empirically known to bind more weakly than the sum of the individual site free energies. This shortfall is generally attributed to the entropy decreases in the linker regions partially offsetting the favorable binding energies.[1,2]

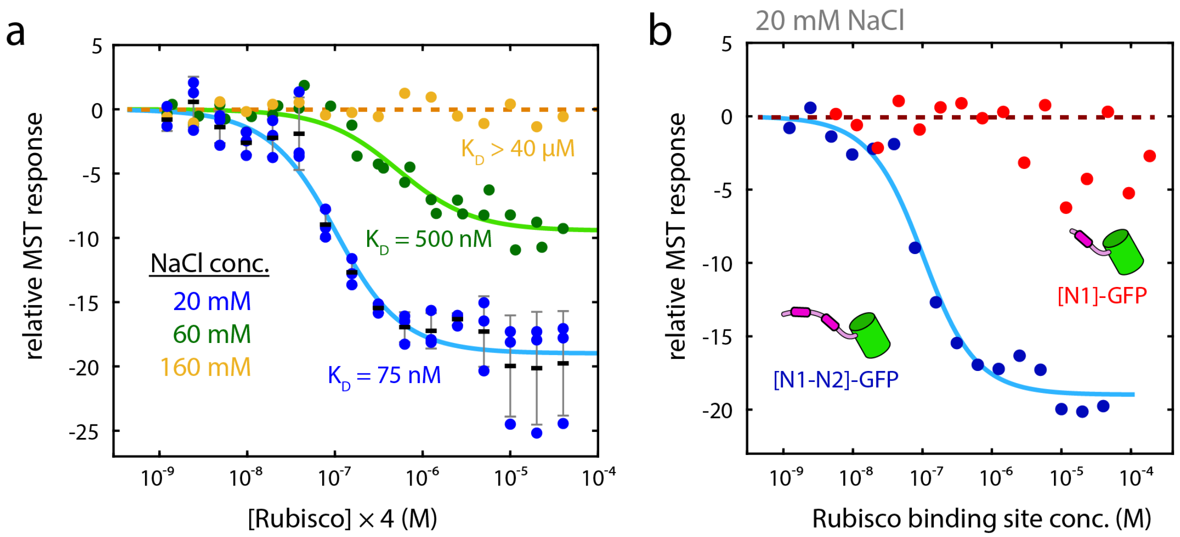

**Figure S5**

**a**, MST responses for [N1-N2]-GFP association to Rubisco. The concentration of the target, [N1-N2]-GFP, was 50 nM. The abscissa represents the concentration of effective binding sites and is four times the Rubisco $L_8S_8$ concentration since each target will engage two of the eight possible sites. Binding experiments were performed at 20, 60, and 160 mM NaCl. At 20 mM NaCl three replicates were performed across 16 Rubisco concentrations. Black lines indicate the means while the gray whiskers show +/- one standard deviation. At 60 mM NaCl the experiment was performed twice with slightly varying concentrations. At 160 mM NaCl data from one representative experiment is shown. The fits to the 20 mM and 60 mM NaCl data are according to Eq. S2 and represent the mean fit parameters from bootstrap sampling of the data. For 160 mM NaCl no binding could be determined over this concentration range and the dashed orange line is drawn at zero response as a visual guide. **b**, Comparison between a double N-peptide, [N1-N2]-GFP, and single N-peptide, [N1]-GFP, species by MST. Both had 50 nM target. The Rubisco binding site concentration is specific to the two different targets. For [N1-N2]-GFP it is the concentration of $L_8S_8$ multiplied by 4 and for [N1]-GFP it is the concentration of $L_8S_8$ multiplied by 8 since the former has four potential binding sites on the Rubisco holoenzyme while the latter has eight. The [N1]-GFP data points are the mean values values from (**a**). The [N1]-GFP data points are from one representative experiment and indicates no conclusive binding over the concentration range. The dashed red line is at zero response as a visual guide.

*Rubisco-N* fusion characterization*

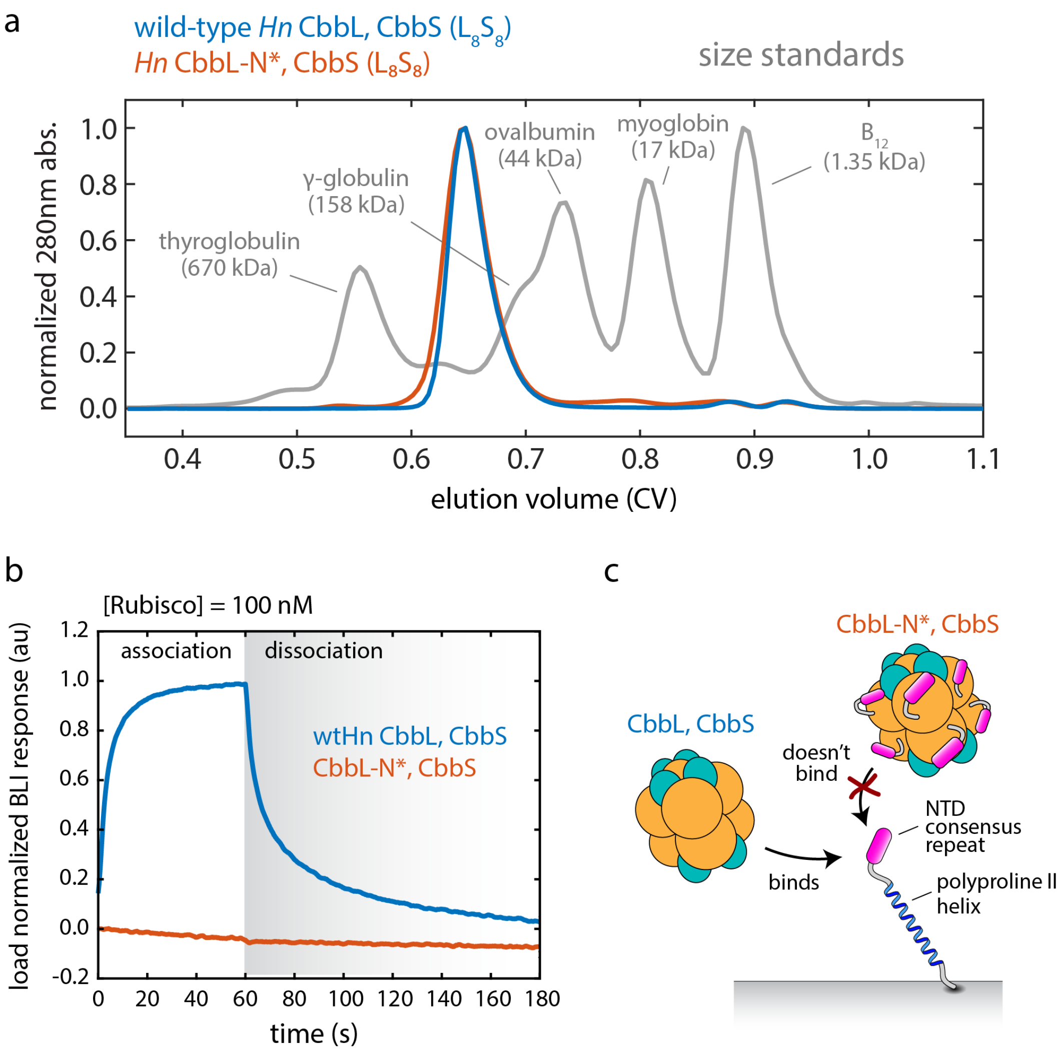

**Figure S6**

**a**, Size exclusion analysis of wild-type Rubisco and the N*-peptide fusion construct. Both elute at volumes commensurate with compact $L_8S_8$ complexes. A run with the Bio-Rad Gel Filtration Standard is included for comparison. Standard masses are indicated. **b**, BLI responses of wtRubisco and the N*-peptide fusion Rubisco at 100 nM with N*-polyPro as the surface bait. The fusion showed no binding. **c**, Proposed cartoon model of differential BLI binding activities. N*-peptide fusion Rubisco is apparently self-passivated by saturating the binding sites from stable association of the fused N*-peptides.

*X-ray crystallography refinement statistics*

Table S1. Data collection and refinement statistics (molecular replacement)

| | *H. neapolitanus* CbbL-N*, CbbS |
|---|---|
| **Data collection** | |
| Space group | C 2 |
| Cell dimensions | |
| $a$, $b$, $c$ (Å) | 171.83 153.95 108.06 |
| $\alpha$, $\beta$, $\gamma$ (°) | 90 124.70 90 |
| Resolution (Å) | 104.1 - 2.4 (2.486 - 2.4)[a] |
| $R_{sym}$ or $R_{merge}$ | 0.1244 (0.5876) |
| $I / \sigma I$ | 12.84 (2.97) |
| Completeness (%) | 99.90 (99.92) |
| Redundancy | 6.9 (6.4) |
| **Refinement** | |
| Resolution (Å) | 104.1 - 2.4 (2.486 - 2.4) |
| No. reflections | 89958 (8958) |
| $R_{work}$ / $R_{free}$ | 0.184 (0.235) / 0.248 (0.301) |
| No. atoms | 18568 |
| Protein | 17618 |
| Ligand/ion | 0 |
| Water | 950 |
| $B$-factors | 41.56 |
| Protein | 41.65 |
| Ligand/ion | n/a |
| Water | 40.04 |
| R.m.s. deviations | |
| Bond lengths (Å) | 0.008 |
| Bond angles (°) | 1.19 |

Values in parentheses are for highest-resolution shell.

a) data from a single crystal

*Enrichment of binding motif from peptide array data*

Cai et al. performed an experiment testing the binding of Rubisco to a peptide array chip.[3] The peptide library was composed of every 8-mer CsoS2 peptide (from *Prochlorococcus marinus* MIT9313) tiled residue-by-residue across the entire protein. The chip was incubated with Rubisco, washed, and then assayed with fluorescently labeled anti-RbcL antibody. The relative fluorescence at each site then provides some indication of the Rubisco binding activity. Cai et al. observed a large number of potential hits scattered throughout the CsoS2 sequence making a determination of the interaction motif challenging. The raw data was generously provided by the authors in the Supplementary Material. This data set was re-analyzed in light of the biochemical and structural evidence of the binding motif presented herein.

Since the original data did not have a clear indication of a specific binding site, we chose to look at it in a statistical manner. R3, R9, and R10 are key conserved N*-peptide residues containable within an 8-mer (G17 and K18 are too far separated). We examined those peptides containing at least two basic residues (i.e. K or R) as generically positively charged species and ones matching the particular R spacings consistent with any pair of arginines among R3, R9, and R10 as a test of the specific motif (i.e. RxxxxxR, RxxxxxxR, or RR).

The results showed that the doubly basic peptides demonstrate enriched fluorescence signal relative to the full peptide library. Peptides matching the motif regular expressions above, however, had significantly greater enrichment even than the doubly basic subset (Fig. S7a). A bootstrap analysis was performed to assess the likelihood of obtaining an equivalently high median fluorescence via random peptides from either the whole population or the doubly basic subset (Fig. S7b). Out of 10,000 trials none were found to exceed the motif median fluorescence, implying a p-value less than $10^{-4}$.

This outcome indicates that the peptide array binding propensities are indeed consistent with the binding motif identified in this work. The relatively high abundance of 8-mers in the peptide library containing portions of the binding motif resulted in multiple “hotspots” scattered throughout the CsoS2 sequence and made prospective unique identification of the specific binding site(s) impossible. The statistical strength of the retrospective motif analysis attests the genuine signal in the peptide array experiment.

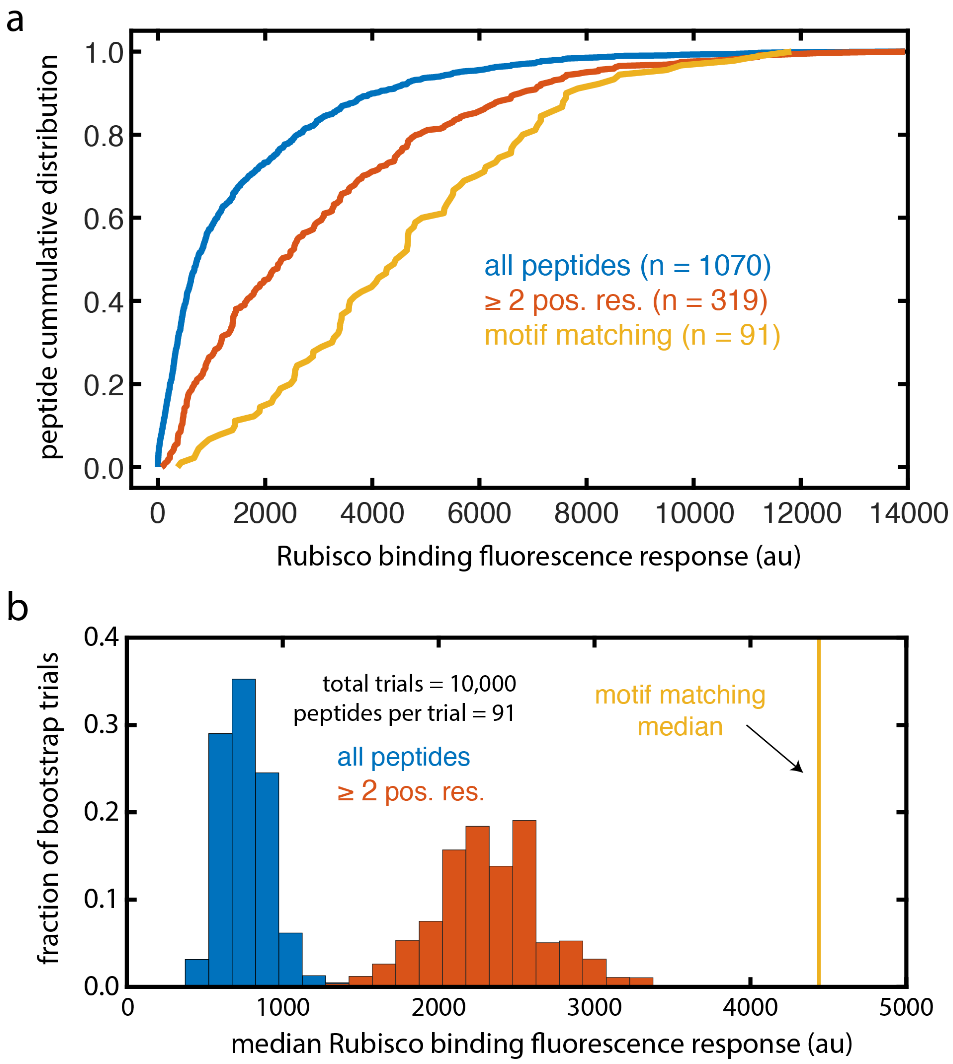

**Figure S7**

**a**, Cumulative distributions of Rubisco binding fluorescence response for CsoS2 array peptides including the full dataset, those with more than two basic residues, and those matching the N*-peptide arginine motif. **b**, Distributions of bootstrap results. 91 peptides were taken at random (with replacement) from either the full dataset or those with two or more basic residues and the median fluorescence response calculated. 10,000 trials were conducted with each set and none exceeded the motif matching median.

*Hydrogen / deuterium exchange of Rubisco inside and outside carboxysomes*

Hydrogen / deuterium exchange (HDX) mass spectrometry experiments were performed on purified Rubisco and Rubisco encapsulated within purified carboxysomes. Overall, the differences between these two states was very minor with no particular regions possessing systematic differential protection (Fig. S8).

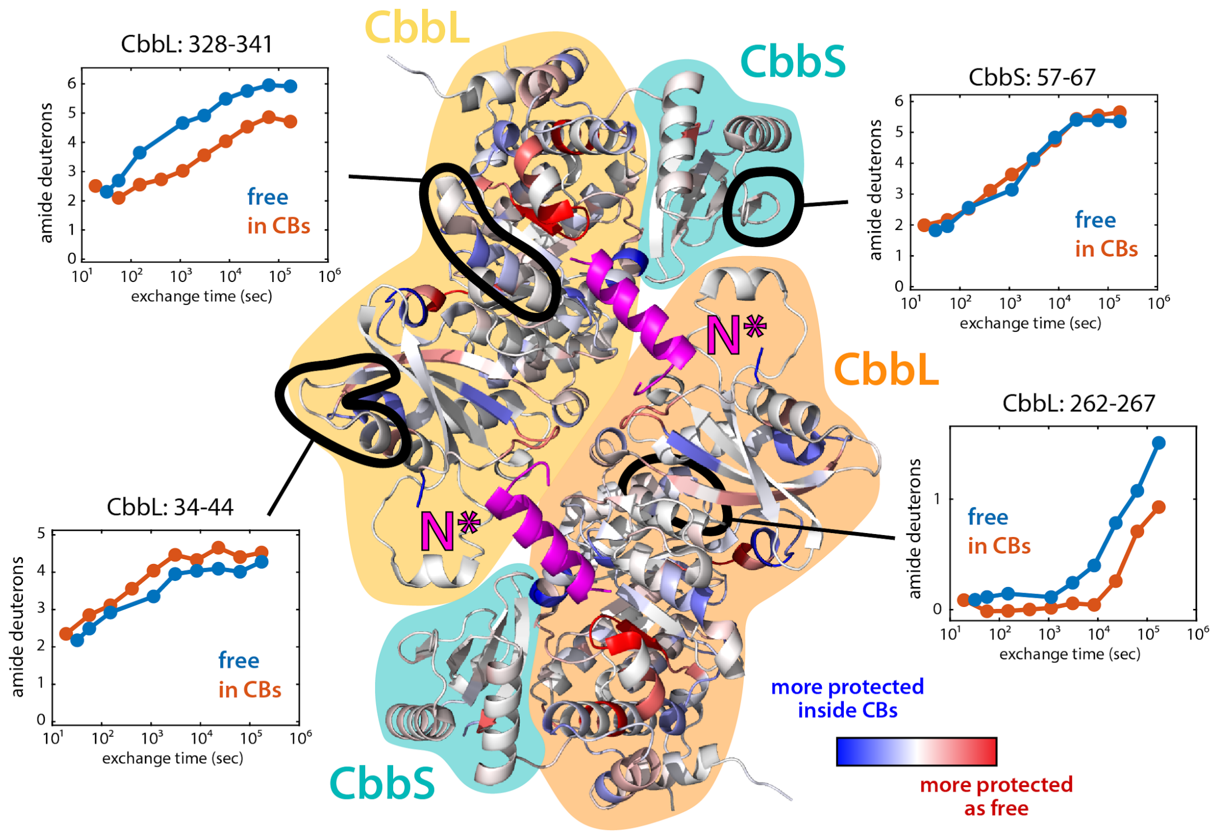

**Figure S8**

The structure displayed contains two CbbLs and two CbbS and shows the $L_2$ dimer interface across which the N*-peptide (in magenta) binds. The Rubisco cartoon is colored according to the differential protection to amide hydrogen exchange. Those residues in blue experience greater protection within purified carboxysomes and those in red experience greater protection as free Rubisco. The comparison between these states was carried out with HDExaminer (Sierra Analytics) using moderate smoothing. Four specific peptides outlined in black highlight some of the diversity of HDX behavior. Most peptides that were observed from both states had essentially identical exchange kinetics as exemplified by the top right subpanel for CbbS: 57-67. Less common were peptides with different exchange profiles between encapsulated and unencapsulated Rubisco. CbbL: 34-44 (lower left subpanel) had slightly more protection in free Rubisco. CbbL: 328-341 (upper left subpanel) and CbbL: 262-267 (lower right subpanel) both had greater protection inside carboxysomes. Since it has the most dramatic results and comes closest to the N*-peptide, the interactions of CbbL: 328-341 are examined in greater detail in Fig. S9 below.

Several specific peptides present in both samples had distinctive exchange profiles. Of particular note is CbbL: 328-341 which showed greater protection inside carboxysomes and is found in relatively close proximity to the N*-peptide (Fig. S8). N* contacts CbbL D340 through a water-mediated hydrogen bond network extending to N* G1 (Fig. S9). The Rubisco-N* fusion protein used for the crystal structure has a two serine linker joining N* to the CbbL C-terminus which is not observed to be ordered. It is possible that residues upstream of the N-peptide binding motifs of CsoS2 interact more extensively with CbbL: 328-341 in the carboxysome. Finally, CbbL: 328-341 covers much of helix 6 and part of loop 6 which plays a role in Rubisco's activity and $CO_2/O_2$ specificity.[4] Consequently, it is conceivable that CsoS2 binding leads to changes in Rubisco's catalytic properties tailored to the unique chemical environment of carboxysome lumen.

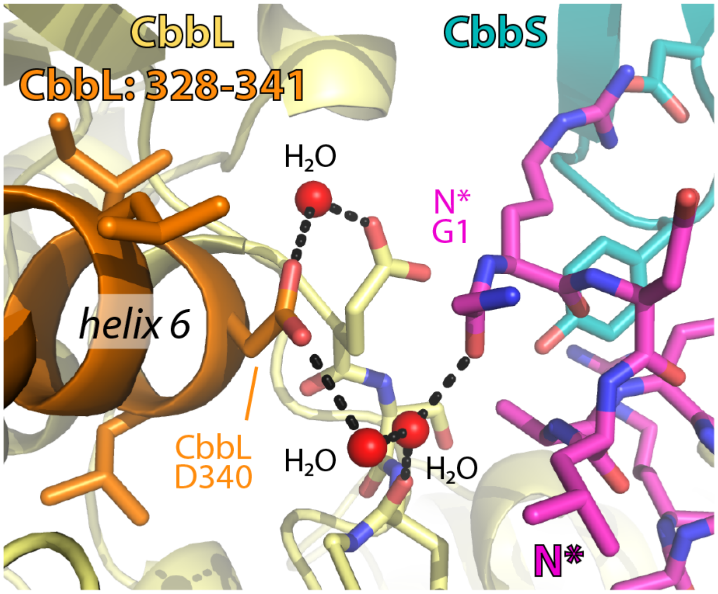

**Figure S9**

Structure of the interactions bridging the N*-peptide (magenta) and CbbL: 328-341 (orange), a peptide demonstrating significant HDX differences between carboxysomal and free Rubisco. A water-mediated hydrogen bond network extends between CbbL D340 and N* G1 perhaps accounting in part for the peptide's greater HDX protection in carboxysomes.

*Structural comparison to CcmM/Rubisco interaction*

The recent work by Wang et al.[5] on the structure of the CcmM/Rubsico complex and attendant liquid-liquid phase separation affords a direct comparison between that scaffold interaction underlying the β-carboxysome assembly and the CsoS2/Rubisco interaction, described herein, central to α-carboxysome assembly. Striking parallels are evident but the molecular details are distinct and bear no obvious evolutionary connection. That both systems converged upon multivalent binding to nearly identical Rubisco sites and have propensities toward phase separation, is a fascinating coincidence and perhaps a hint at some optimality of this assembly strategy.

In both cases the scaffolding element binds at the union of two $L_2$ dimers and a small subunit (Fig. S10a). Consequently, the binding site only exists in the fully assembled $L_8S_8$ Rubisco holoenzyme. Unlike the N*-peptide which can apparently simultaneously bind at eight possible sites, CcmM-SSUL occludes the immediately adjacent site and therefore has only four possible sites per Rubisco.

Wang et al. identified two primary regions of interaction between CcmM-SSUL and Rubisco which they called “Interface I” and “Interface II”. Interface I closely overlaps with the N*/Rubisco binding and is the area of focus below in Fig. S10b. (Interface II is located near the loop at the bottom right and is not shown in detail.) Interface I is largely electrostatic in nature with a series of three arginines (CcmM R251, R252, and R254) making important contacts. R251 and R252 form salt bridges to two apartates (RbcS D93 and RbcL D76) which are positionally equivalent to those engaged in salt bridges to N* R2 and N* R10, respectively (see Fig. S10b table). Despite utilizing some of the same residues for salt bridges, the scaffold geometries are remarkably different. In CcmM, residues R251, R252, and R254 fan out from a short helix, called *α2*, whose axis is directed down inward to Rubisco. The N*-peptide’s helix axis, in contrast, runs perpendicular to the Rubisco surface. Finally, CcmM/Rubisco contains no apparent cation-π interactions which feature prominently in the N*/Rubisco binding interface.

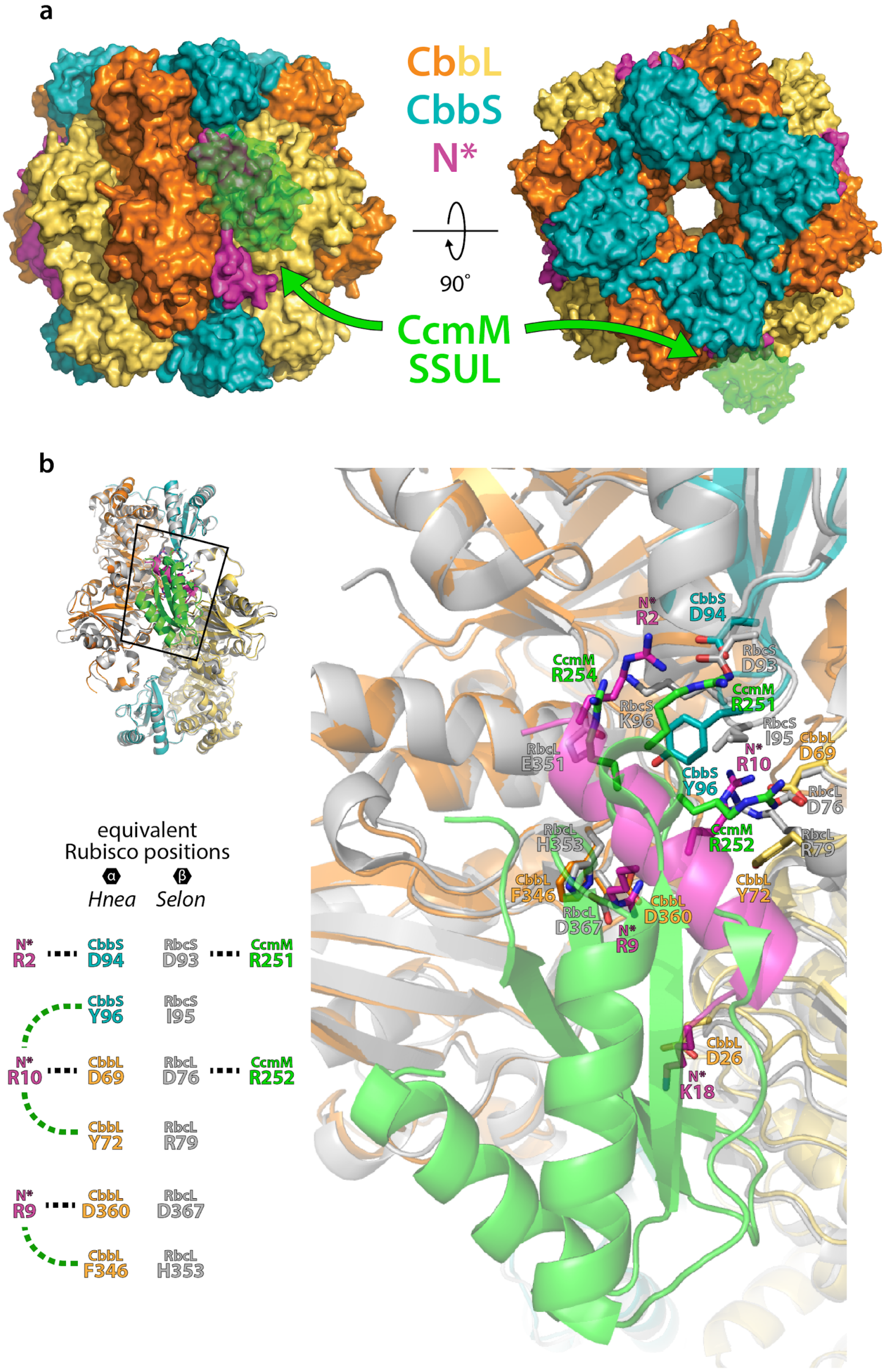
a
CbbL
CbbS
N*
90˚
CcmM
SSUL
b
equivalent
Rubisco positions
α
β
Hnea
Selon
N* R2
CbbS D94
RbcS D93
CcmM R251
CbbS Y96
RbcS I95
N* R10
CbbL D69
RbcL D76
CcmM R252
CbbL Y72
RbcL R79
N* R9
CbbL D360
RbcL D367
CbbL F346
RbcL H353
CcmM R254
RbcS K96
RbcL E351
CbbL D26
N* K18

**Figure S10**

**a**, Surface representation of the N*/Rubisco complex with aligned CcmM-SSUL from the model of Wang et al.[5] in semi-transparent green. **b**, Detailed comparative view of the scaffold/Rubisco interaction interface. The inset table pairs equivalent Rubisco positions from alignment and the dashed lines indicate select specific interactions to the corresponding scaffold element shown with salt bridges in black and cation-π interactions in green. “*Hnea*” is the α-carboxysomal Form IA Rubisco from *Halothiobacillus neapolitanus* with CbbL (in orange/yellow) and CbbS (in cyan). The N*-peptide-bound structure is from the current study with PDB ID: XXXX. “*Selon*” is the β-carboxysomal Form IB Rubisco from *Synechococcus elongatus PCC 7942* with large subunit, RbcL, and small subunit, RbcS, both in grey. The bound small subunit-like repeat, CcmM-SSUL1, is shown in green. The atomic model was determined from cryo-electron microscopy single particle analysis and has PDB ID: 6HBC.

*CsoS2 NTD truncations and carboxysome formation*

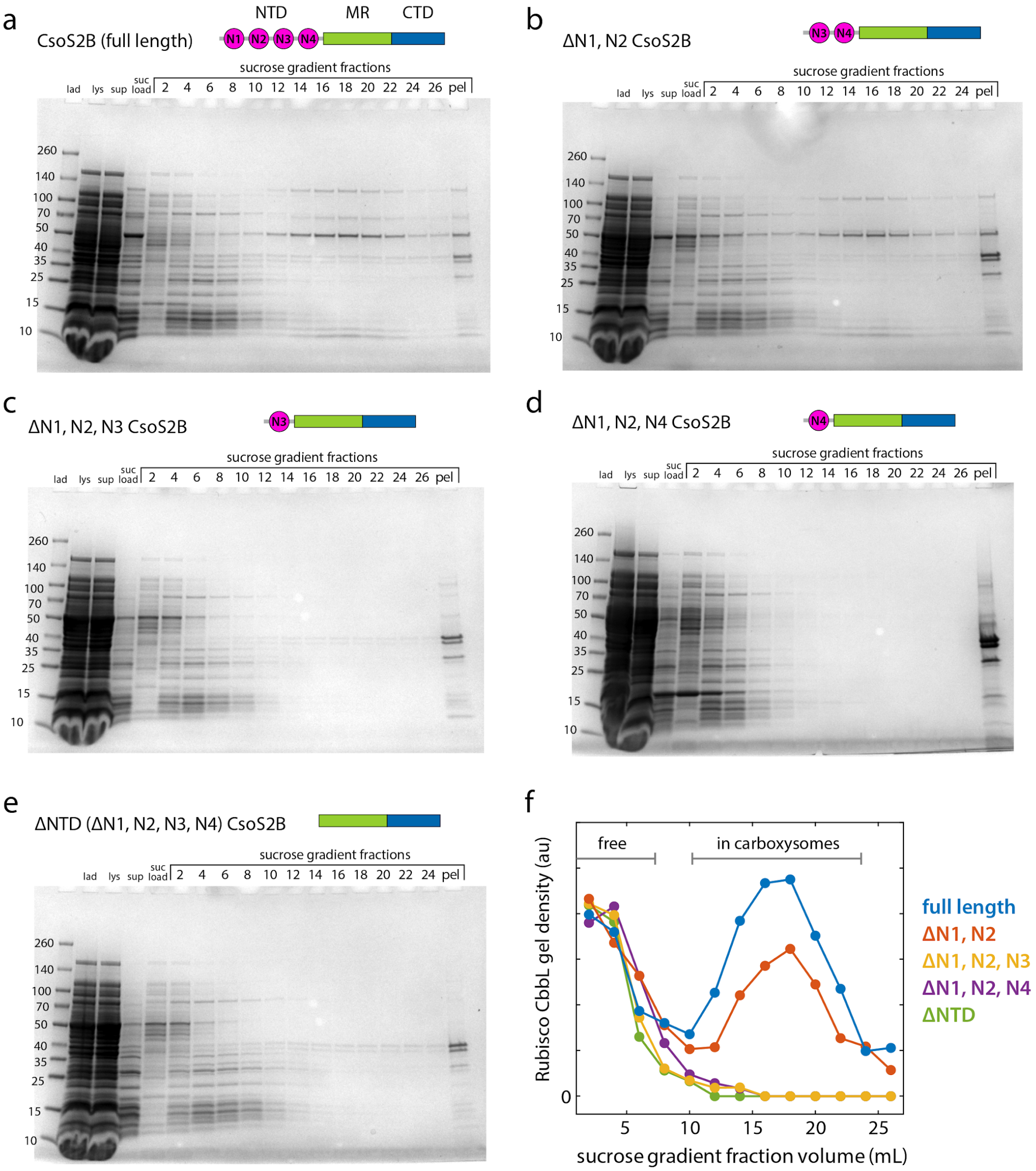

**Figure S11**

**a-e**, 4-20% SDS-PAGE gels of carboxysome purifications for each of the NTD truncation constructs, shown schematically. The carboxysomes were purified according to established protocols culminating with ultracentrifugation on a sucrose step gradient having 5-mL layers of 10, 20, 30, 40, and 50% (w/v) sucrose. Each fraction was 1 mL. The “pel” fraction is the resuspended pellet from bottom of the gradient. Normal carboxysomes typically occur as a broad band peaked around 18 mL. **f**, Rubisco large subunit gel density as a function of fraction volume. As the major carboxysome component, Rubisco is a sensitive proxy for the intact particles. Only full length CsoS2 and one retaining two of the N-peptide repeats (ΔN1, N2) resulted in purifiable carboxysomes.

*Protein sequences*

Select features and mutation sites are indicated by highlights and described after each sequence.

***Halothiobacillus neapolitanus* Rubisco:**

CbbL (large subunit):

MSAVKKYSAGVKEYRQTYWMPEYTPLDSDILACFKITPQPGVDREEAAAAVAAESSTGTWTTV
WTDLLTDMDYYKGRAYRIEDVPGDDAAFYAFIAYPIDLFEEGSVVNVFTSLVGNVFGFKAVRGL
RLEDVRFPLAYVKTCGGPPHGIQVERDKMNKYGRPLLGCTIKPKLGLSAKNYGRAVYECLRGG
LDFTKDDENINSQPFMRWRDRFLFVQDATETAEAQTGERKGHYLNVTAPTPEEMYKRAEFAK
EIGAPIIMHDYITGGFTANTGLAKWCQDNGVLLHIHRAMHAVIDRNPNHGIHFRVLTKILRLSGG
DHLHTGTVVGKLEGDRASTLGWIDLLRESFIPEDRSRGIFFDQDWGSMPGVFAVASGGIHVWH
MPALVNIFGDDSVLQFGGGTLGHPWGNAAGAAANRVALEACVEARNQGRDIEKEGKEILTAAA
QHSPELKIAMETWKEIKFEFDTVDKLDTQNRWSHPQFEK

CbbS (small subunit):

MAEMQDYKQSLKYETFSYLPPMNAERIRAQIKYAIAQGWSPGIEHVEVKNSMNQYWYMWKLP
FFGEQNVDNVLAEIEACRSAYPTHQVKLVAYDNYAQSLGLAFVVYRGN

StrepII affinity tag
CbbL Y72; Y72A or Y72R
CbbL F346; F346A
CbbS Y96; Y96A

***Halothiobacillus neapolitanus* CsoS2:**

MHHHHHHPSQSGMNPADLSGLSGKELARARRAALSKQGKAAVSNKTASVNRSTKQAASSINT
NQVRSSVNEVPTDYQMADQLCSTIDHADFGTESNRVRDLCRQRREALSTIGKKAVKTNGKPS
GRVRPQQSVVHNDAMIENAGDTNQSSSTSLNNELSEICSIADDMPERFGSQAKTVRDICRARR
QALSERGTRAVPPKPQSQGGPGRNGYQIDGYLDTALHGRDAAKRHREMLCQYGRGTAPSCK
PTGRVKNSVQSGNAAPKKVETGHTLSGGSVTGTQVDRKSHVTGNEPGTCRAVTGTEYVGTE
QFTSFCNTSPKPNATKVNVTTTARGRPVSGTEVSRTEKVTGNESGVCRNVTGTEYMSNEAHF
SLCGTAAKPSQADKVMFGATARTHQVVSGSDEFRPSSVTGNESGAKRTITGSQYADEGLARL
TINGAPAKVARTHTFAGSDVTGTEIGRSTRVTGDESGSCRSISGTEYLSNEQFQSFCDTKPQR
SPFKVGQDRTNKGQSVTGNLVDRSELVTGNEPGSCSRVTGSQYGQSKICGGGVGKVRSMRT
LRGTSVSGQQLDHAPKMSGDERGGCMPVTGNEYYGREHFEPFCTSTPEPEAQSTEQSLTCE
GQIISGTSVDASDLVTGNEIGEQQLISGDAYVGAQQTGCLPTSPRFNQTGNVQSMGFKNTNQP
EQNFAPGEVMPTDFSIQTPARSAQNRITGNDIAPSGRITGPGMLATGLITGTPEFRHAARELVG
SPQPMAMAMANRNKAAQAPVVQPEVVATQEKPELVCAPRSDQMDRVSGEGKERCHITGDD
WSVNKHITGTAGQWASGRNPSMRGNARVVETSAFANRNVPKPEKPGSKITGSSGNDTQGSLI
TYSGGARG

Hexahistidine affinity tag
NTD peptide repeats
Middle region peptide repeats
CTD peptide repeats
CTP

**CsoS2 N-terminal domain (NTD):**

MSHHHHHHPSQSGMNPADLSGLSGKELARARRAALSKQGKAAVSNKTASVNRSTKQAASSIN
TNQVRSSVNEVPTDYQMADQLCSTIDHADFGTESNRVRDLCRQRREALSTIGKKAVKTNGKPS
GRVRPQQSVVHNDAMIENAGDTNQSSSTSLNNELSEICSIADDMPERFGSQAKTVRDICRARR
QALSERGTRAVPPKPQSQGGPGRNGYQIDGYLDTALHGRDAAKRHREMLCQYGRGTAPSCK
PTGRVKNSVQSGNAAPKKV

Hexahistidine affinity tag
NTD peptide repeats
Basic residues in N-peptides all mutated to alanines

**CsoS2 Middle Region (MR):**

MSHHHHHHAPKKVETGHTLSGGSVTGTQVDRKSHVTGNEPGTCRAVTGTEYVGTEQFTSFCNTSPKPNATKVNVTTTARGRPVSGTEVSRTEKVTGNESGVCRNVTGTEYMSNEAHFSLCGTAAKPSQADKVMFGATARTHQVVSGSDEFRPSSVTGNESGAKRTITGSQYADEGLARLTINGAPAKVARTHTFAGSDVTGTEIGRSTRVTGDESGSCRSISGTEYLSNEQFQSFCDTKPQRSPFKVGQDRTNKGQSVTGNLVDRSELVTGNEPGSCSRVTGSQYGQSKICGGGVGKVRSMRTLRGTSVSGQQLDHAPKMSGDERGGCMPVTGNEYYGREHFEPFCTSTPEPEAQ

Hexahistidine affinity tag
Middle region peptide repeats

**CsoS2 C-terminal domain (CTD):**

MSHHHHHHTSTPEPEAQSTEQSLTCEGQIISGTSVDASDLVTGNEIGEQQLISGDAYVGAQQTGCLPTSPRFNQTGNVQSMGFKNTNQPEQNFAPGEVMPTDFSIQTPARSAQNRITGNDIAPSGRITGPGMLATGLITGTPEFRHAARELVGSPQPMAMAMANRNKAAQAPVVQPEVVATQEKPELVCAPRSDQMDRVSGEGKERCHITGDDWSVNKHITGTAGQWASGRNPSMRGNARVVETSAFANRNVPKPEKPGSKITGSSGNDTQGSLITYSGGARG

Hexahistidine affinity tag
CTD peptide repeats
CTP

**N*-polyProline:**

MSWKHHHHHHENLYFQSAAVGGGSGGGSGGPPPPPPPPPAPAPAPAPPPPPPPPPAPAPAPAPPPPPPPPGGGSGGGSGGGRDLARARREALSQQGKAAVGGGSGGGSGGSG

Hexahistidine affinity tag
polyproline II helix
NTD peptide consensus repeat (N*)

**random N*-polyProline:**

MSWKHHHHHHENLYFQSAAVGGGSGGGSGGPPPPPPPPPAPAPAPAPPPPPPPPPAPAPAPAPPPPPPPPGGGSGGGSGGGRRKGLRAAGRALQVEQADSRAGGGSGGGSGGSG

Hexahistidine affinity tag
polyproline II helix
random NTD peptide

**polyProline flexible polar AA control:**

MSWKHHHHHHENLYFQSAAVGGGSGGGSGGPPPPPPPPPAPAPAPAPPPPPPPPPPAPAPAPAPPPPPPPPGGGSGGGSGGGGTGSTGSGSSSSGSGTSGTGGGGSGGGSGGSG

Hexahistidine affinity tag

polyproline II helix

**MR R1-polyProline:**

MSWKHHHHHHENLYFQSAAVGGGSGGGSGGPPPPPPPPPAPAPAPAPPPPPPPPPPAPAPAPAPPPPPPPPGGGSGGGSGGGKVETGHTLSGGSVTGTQVDRKSHVTGNEPGTCRAVTGTEYVGTEQFTSFCGGGSGGGSGGSG

Hexahistidine affinity tag

polyproline II helix

MR repeat 1

**CTD R8-polyProline:**

MSWKHHHHHHENLYFQSAAVGGGSGGGSGGPPPPPPPPPAPAPAPAPPPPPPPPPPAPAPAPAPPPPPPPPGGGSGGGSGGGSIQTPARSAQNRITGNDIAPSGRITGPGMLATGLITGTPEFGGGSGGGSGGSG

Hexahistidine affinity tag

polyproline II helix

CTD repeat 8

**CTP-polyProline:**

MSWKHHHHHHENLYFQSAAVGGGSGGGSGGPPPPPPPPPAPAPAPAPPPPPPPPPPAPAPAPAPPPPPPPPGGGSGGGSGGGVPKPEKPGSKITGSSGNDTQGSLITYSGGARGGGGSGGGSGGSG

Hexahistidine affinity tag

polyproline II helix

CTP

**[N1-N2]-GFP:**

MHHHHHHENLYFQSPSQSGMNPADLSGLSGKELARARRAALSKQGKAAVSNKTASVNRSTK
QAASSINTNQVRSSVNEVPTDYQMADQLCSTIDHADFGTESNRVRDLCRQRREALSTIGKKAV
KTNGKPSGRVRPQQSGSGSGGSKGEELFTGVVPILVELDGDVNGHKFSVRGEGEGDATNGK
LTLKFICTTGKLPVPWPTLVTTLTYGVQCFSRYPDHMKQHDFFKSAMPEGYVQERTISFKDDGT
YKTRAEVKFEGDTLVNRIELKGIDFKEDGNILGHKLEYNFNSHNVYITADKQKNGIKANFKIRHNV
EDGSVQLADHYQQNTPIGDGPVLLPDNHYLSTQSKLSKDPNEKRDHMVLLEFVTAAGITHGMD
ELYKWSHPQFEK

Hexahistidine affinity tag
NTD peptide repeats (N1 and N2)
NTD flexible “interstitial” sequence
Superfolder GFP [6] (with dimer abolishing K206 variant) [7]

**[N1]-GFP:**

MSHHHHHHPSQSGMNPADLSGLSGKELARARRAALSKQGKAAVSNKTASVNRSTKQAASSIN
TNQVRSSVNEVPTDYQMADQLCSTIDHADFGTESNRVENLYFQSSGSGSGGSKGEELFTGVV
PILVELDGDVNGHKFSVRGEGEGDATNGKLTLKFICTTGKLPVPWPTLVTTLTYGVQCFSRYPD
HMKQHDFFKSAMPEGYVQERTISFKDDGTYKTRAEVKFEGDTLVNRIELKGIDFKEDGNILGHK
LEYNFNSHNVYITADKQKNGIKANFKIRHNVEDGSVQLADHYQQNTPIGDGPVLLPDNHYLSTQ
SKLSKDPNEKRDHMVLLEFVTAAGITHGMDELYKWSHPQFEK

Hexahistidine affinity tag
NTD peptide repeat (N1)
NTD flexible “interstitial” sequence
Superfolder GFP (with dimer abolishing K206 variant)

**Rubisco-N* fusion (CbbL-N*, CbbS):**

CbbL-N*:

MSAVKKYSAGVKEYRQTYWMPEYTPLDSDILACFKITPQPGVDREEAAAAVAAESSTGTWTTVWTDLLTDMDYYKGRAYRIEDVPGDDAAFYAFIAYPIDLFEEGSVVNVFTSLVGNVFGFKAVRGLRLEDVRFPLAYVKTCGGPPHGIQVERDKMNKYGRPLLGCTIKPKLGLSAKNYGRAVYECLRGGLDFTKDDENINSQPFMRWRDRFLFVQDATETAEAQTGERKGHYLNVTAPTPEEMYKRAEFAKEIGAPIIMHDYITGGFTANTGLAKWCQDNGVLLHIHRAMHAVIDRNPNHGIHFRVLTKILRLSGGDHLHTGTVVGKLEGDRASTLGWIDLLRESFIPEDRSRGIFFDQDWGSMPGVFAVASGGIHVWHMPALVNIFGDDSVLQFGGGTLGHPWGNAAGAAANRVALEACVEARNQGRDIEKEGKEILTAAAQHSPELKIAMETWKEIKFEFDTVDKLDTQNRSSGRDLARARREALSQQGKAAVGSWSHPQFEK

CbbS:

MAEMQDYKQSLKYETFSYLPPMNAERIRAQIKYAIAQGWSPGIEHVEVKNSMNQYWYMWKLPFFGEQNVDNVLAEIEACRSAYPTHQVKLVAYDNYAQSLGLAFVVYRGN

StrepII affinity tag

NTD consensus repeat

*Plasmids used*

All plasmids in the table below were made for this study.

Table S2

| ***Name*** | ***Relevant genotype*** | ***Resistance*** |
|---|---|---|
| pBz15 | wild-type *Halothiobacillus neapolitanus* Form I Rubisco with StrepII tag (CbbL-StrepII, CbbS), pET-14b backbone | Amp |
| pLz74 | Same as pBz15 but with N*-peptide fusion on CbbL, construct used for protein crystallization | Amp |
| pBz118 | His-tagged *Hn* CsoS2. This protein contains a programmed ribosomal frameshift and expresses as a short and long form (CsoS2A and CsoS2B, respectively),[8] pET-14b backbone | Amp |
| pBz109 | His-tagged CsoS2 N-terminal domain (NTD), pET-14b backbone | Amp |
| pBz106 | His-tagged CsoS2 Middle Region (MR), pET-14b backbone | Amp |
| pBz110 | His-tagged CsoS2 C-terminal domain (CTD), pET-14b backbone | Amp |
| pLz47 | Consensus N*-peptide after polyproline II helix, N-terminal His-tag, pET-14b backbone | Amp |
| pLz55 | Same as pLz47 but with randomized N*-peptide sequence | Amp |
| pLz26 | His-tagged first two N-peptides of NTD fused to Superfolder GFP ([N1-N2]-GFP) used for MST, pET-14b backbone | Amp |
| pBz172 | His-tagged first N-peptide of NTD fused to Superfolder GFP ([N1]-GFP) used for MST, pET-14b backbone | Amp |
| pLz37 | Derived from pHnCB10 for heterologous expression of carboxysome operon proteins.[9] Contains full-length CsoS2. | Cm |
| pLz75 | Same as pLz37 but with CsoS2 truncated such that NTD repeats N1 and N2 are removed | Cm |
| pLz76 | Same as pLz37 but with CsoS2 truncated such that NTD repeats N1, N2, and N3 are removed | Cm |
| pLz77 | Same as pLz37 but with CsoS2 discontinuously truncated such that repeats N1, N2, and N4 are removed | Cm |
| pLz78 | Same as pLz37 but with CsoS2 truncated to remove the entire NTD (i.e. N1, N2, N3, and N4 are removed) | Cm |
| pLz54 | Same as pLz47 but with 20 polar amino acids instead of N*-peptide | Amp |

| | | |
|---|---|---|
| pLz56 | Same as pLz47 but with MR repeat 1 instead of N*-peptide | Amp |
| pLz57 | Same as pLz47 but with CTD repeat 8 instead of N*-peptide | Amp |
| pLz58 | Same as pLz47 but with C-terminal peptide (CTP) instead of N*-peptide | Amp |

### References

1. Kane, R. S. Thermodynamics of multivalent interactions: influence of the linker. *Langmuir* **26,** 8636–8640 (2010).
2. Kitov, P. I. & Bundle, D. R. On the nature of the multivalency effect: a thermodynamic model. *J. Am. Chem. Soc.* **125,** 16271–16284 (2003).
3. Cai, F. *et al.* Advances in understanding carboxysome assembly in prochlorococcus and synechococcus implicate csos2 as a critical component. *Life (Basel)* **5,** 1141–1171 (2015).
4. Karkehabadi, S., Satagopan, S., Taylor, T. C., Spreitzer, R. J. & Andersson, I. Structural analysis of altered large-subunit loop-6/carboxy-terminus interactions that influence catalytic efficiency and CO2/O2 specificity of ribulose-1,5-bisphosphate carboxylase/oxygenase. *Biochemistry* **46,** 11080–11089 (2007).
5. Wang, H. *et al.* Rubisco condensate formation by CcmM in β-carboxysome biogenesis. *Nature* **566,** 131–135 (2019).
6. Pédelacq, J.-D., Cabantous, S., Tran, T., Terwilliger, T. C. & Waldo, G. S. Engineering and characterization of a superfolder green fluorescent protein. *Nat. Biotechnol.* **24,** 79–88 (2006).
7. Zacharias, D. A., Violin, J. D., Newton, A. C. & Tsien, R. Y. Partitioning of lipid-modified monomeric GFPs into membrane microdomains of live cells. *Science* **296,** 913–916 (2002).
8. Chaijarasphong, T. *et al.* Programmed Ribosomal Frameshifting Mediates Expression of the α-Carboxysome. *J. Mol. Biol.* **428,** 153–164 (2016).
9. Bonacci, W. *et al.* Modularity of a carbon-fixing protein organelle. *Proc Natl Acad Sci USA* **109,** 478–483 (2012).
